# A Spatially Coordinated Keratinocyte-Fibroblast Circuit Recruits MMP9^+^ Myeloid Cells to Drive IFN-I-Driven Inflammation in Photosensitive Autoimmunity

**DOI:** 10.1101/2025.08.19.670635

**Authors:** Yuqing Wang, Khashayar Afshari, Nazgol-Sadat Haddadi, Carolina Salomão Lopes, Chee-Huat Linus Eng, Nuria Martinez, Leah Whiteman, Ksenia S Anufrieva, Kevin Wei, Kirsten Frieda, Stefania Gallucci, Misha Rosenbach, Ruth Ann Vleugels, John E Harris, Mehdi Rashighi, Manuel Garber

## Abstract

Photosensitivity is a hallmark of cutaneous lupus erythematosus (CLE) and dermatomyositis (DM), yet the mechanisms linking ultraviolet B (UVB) exposure to tissue-specific autoimmunity remain incompletely defined. Here, we use an integrative human-based approach, including single-cell RNA sequencing, spatial transcriptomics (seqFISH+), in vivo UVB provocation, and in vitro modeling, to uncover a spatially coordinated inflammatory circuit that underlies interferon-I (IFN-I)-amplified skin pathology.

We identify MMP9^+^ CD14^+^ myeloid cells as central effectors of photosensitivity in both CLE and DM. These cells are markedly expanded in lesional skin, serve as the dominant source of IFN-β, and colocalize with cytotoxic CD4^+^ T cells at the dermal–epidermal junction. Spatial transcriptomics further reveals a keratinocyte-fibroblast-myeloid axis, wherein keratinocytes activate discrete subsets of pro-inflammatory fibroblasts in the superficial dermis to produce monocyte-attracting chemokines, including CCL2, CCL19, CCL7, CCL8, and CXCL12, directing MMP9^+^ CD14^+^ cell recruitment toward the interface.

In our in-vitro model, IFN-I-primed basal keratinocytes undergo heightened UVB-induced cell death and release membrane-associated cytokines such as TNF-α, IL-1α, which activate monocyte-derived dendritic cells (moDCs) and induce transcriptional programs mirroring those of MMP9^+^ CD14^+^ cells in vivo. In vivo, UVB irradiation of non-lesional DM skin, but not healthy controls, elicits rapid infiltration of these myeloid cells, confirming their disease-specific responsiveness to UVB.

Finally, in a proof-of-concept clinical study, treatment with anifrolumab (anti-IFN-I receptor) blocked UVB-induced MMP9^+^ CD14^+^ infiltration and attenuated photosensitivity in CLE.

Together, these findings define a multicellular inflammatory cascade linking keratinocyte injury, fibroblast chemotactic programming, and myeloid effector function in IFN-I-driven skin autoimmunity and nominate MMP9^+^ CD14^+^ cells as actionable targets in photosensitive dermatoses.

Photosensitivity is central to cutaneous lupus erythematosus (CLE) and dermatomyositis (DM), but the mechanisms linking UVB exposure to tissue-specific autoimmunity are poorly defined. Using single-cell RNA sequencing, spatial transcriptomics, UVB provocation, and in vitro modeling, we identify MMP9^+^ CD14^+^ myeloid cells as critical mediators of photosensitivity. These cells expand significantly in lesional skin, produce IFN-β, and colocalize with cytotoxic CD4^+^ T cells at the dermal-epidermal junction. Keratinocytes activate fibroblasts in the superficial dermis, prompting them to release chemokines (CCL2, CCL19, CCL7, CCL8, CXCL12) that recruit MMP9^+^ CD14^+^ cells. IFN-I-primed keratinocytes exposed to UVB release cytokines activating dendritic cells, mirroring in vivo responses. UVB irradiation of non-lesional DM skin rapidly recruits these myeloid cells. In a clinical proof-of-concept study, anti-IFN-I treatment with anifrolumab prevented UVB-induced myeloid infiltration and reduced photosensitivity. Thus, targeting MMP9^+^ CD14^+^ cells may offer therapeutic potential for managing photosensitive autoimmune skin conditions.

## Introduction

Ultraviolet (UV) light presents a therapeutic paradox in cutaneous immunology: while broadly immunosuppressive and used to treat diseases such as psoriasis and vitiligo, it exacerbates inflammation in photosensitive dermatoses like cutaneous lupus erythematosus (CLE) and dermatomyositis (DM). The mechanisms underlying this divergent response remain poorly defined^1^.

CLE and DM are characterized by interface dermatitis and enriched type I interferon (IFN-I) signatures, yet how UVB irradiation leads to sustained inflammation in these contexts is unclear. A prevailing model suggests that UV-induced keratinocyte death releases nucleic acids, triggering IFN production^2^. However, this fails to explain subsequent inflammatory responses, or their cellular drivers differ between photosensitive and photoresponsive diseases.

To address this, we employed an integrative multiomics approach across four inflammatory skin conditions, two photosensitive diseases (CLE, DM), and two photoresponsive diseases (psoriasis and vitiligo). By combining single-cell RNA sequencing (scRNA-seq), spatial transcriptomics (seqFISH^3^), and in vivo UVB provocation, we sought to delineate disease-specific immune architecture and identify key effector pathways that mechanistically link UVB exposure to skin inflammation.

Strikingly, the most pronounced disease-specific divergence was observed within the myeloid compartment. While psoriasis and vitiligo lesions contained predominantly conventional dendritic cell subsets, photosensitive lesions were distinguished by an expansion of type I interferon-producing myeloid populations, plasmacytoid dendritic cells in CLE, and CD14^+^ myeloid cells in DM and, to a lesser extent CLE. These CD14^+^ cells, largely overlooked in the context of UV-driven autoimmunity, emerged as dominant IFN-β producers and were spatially enriched at the dermal-epidermal junction, co-localizing with cytotoxic CD4^+^ T cells.

Here, we show that MMP9^+^ CD14^+^ myeloid cells orchestrate UVB-induced inflammation by linking basal keratinocyte damage to T cell–mediated interface dermatitis via inflammatory fibroblasts and myeloid cells. Using in vitro and in vivo human models, we demonstrate that IFN-β–primed keratinocytes are more susceptible to UVB-induced death and produce inflammatory mediators that recruit and activate inflammatory myeloid cells. These, in turn, amplify inflammation via IFN-β production and chemokine secretion, forming a pathogenic feedback loop. Our findings uncover a critical mechanism of photosensitivity and nominate MMP9^+^ CD14^+^ cells as potential therapeutic targets in CLE and DM.

## Results

To define shared and disease-specific immune features in photosensitive versus photoresponsive skin conditions, we performed single-cell RNA sequencing (scRNA-seq) and proteomic profiling on suction blister biopsies from healthy controls as well as lesional and non-lesional skin of patients with DM and CLE (photosensitive) and psoriasis and vitiligo (photoresponsive), ( study subjects information in Table S1), using methods we previously established^4^. Interstitial fluid from each blister was simultaneously analyzed by multiplex immunoassay to quantify 250 soluble proteins^5^ (Fig. 1A and Table S2).

**Figure 1.**
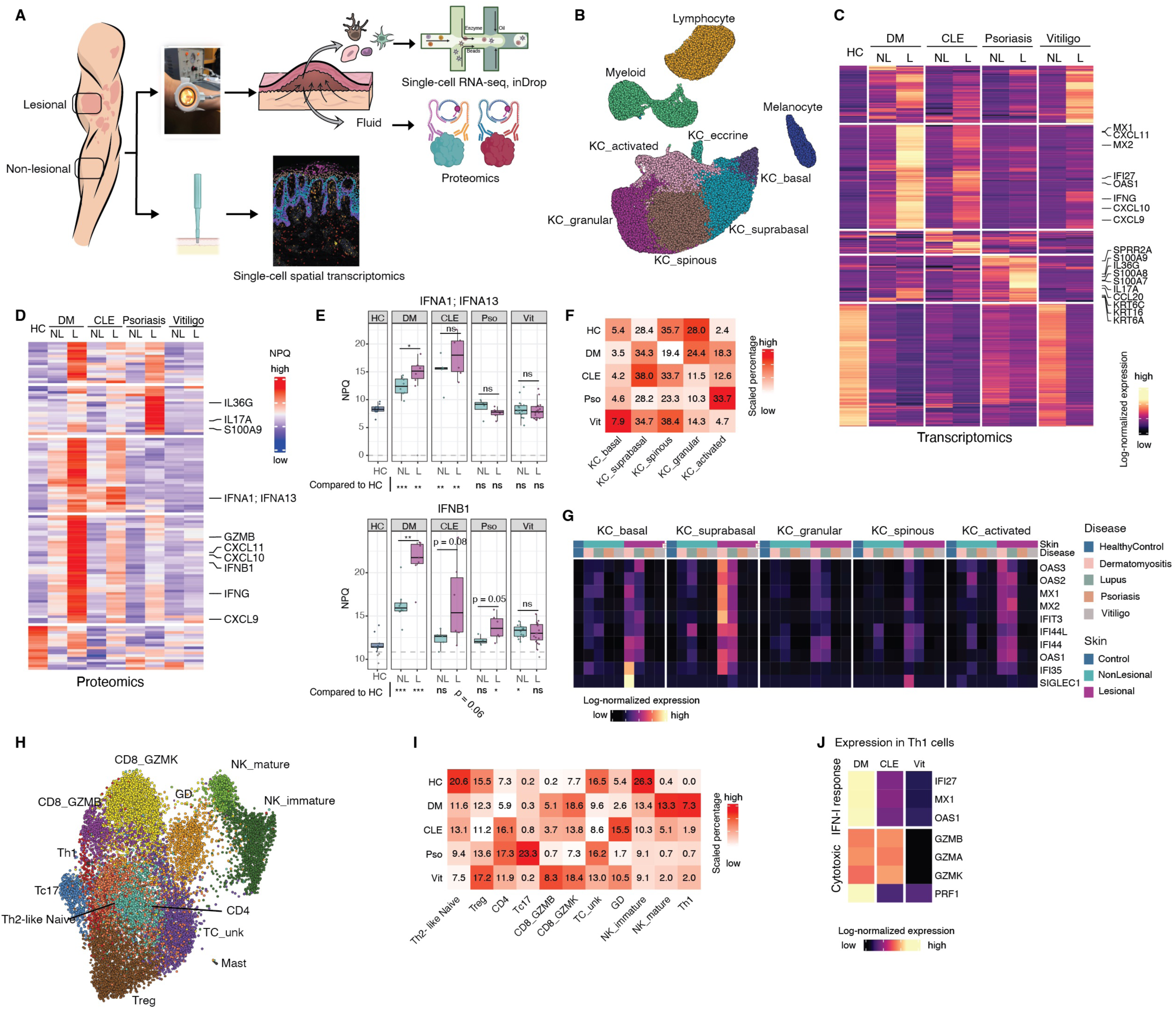
Integrated single-cell transcriptomic and proteomic profiling reveals IFN-I–associated immune and epithelial features across photosensitive and photoresponsive skin diseases. **(A)** Schematic overview of the study. Suction blister biopsies were collected from lesional (L) and non-lesional (NL) skin of patients with dermatomyositis (DM), cutaneous lupus erythematosus (CLE), psoriasis (Pso), vitiligo (Vit), and from healthy controls (HC). Cells in the blister biopsies were used for inDrop scRNA-seq, and the blister fluid was used for proteomics NUcleic acid Linked Immuno-Sandwich Assay (NULISA™). Punch biopsies were collected for spatial transcriptomics sequential Fluorescence In Situ Hybridization (seqFISH). **(B)** UMAP embedding of all cells from blister biopsy scRNA-seq. Cells are colored by cell type. **(C)** Heatmap of differentially expressed genes from pseudo bulk analysis across conditions in scRNA-seq. The top 100 significant genes per disease by fold change (FDR < 0.01), identified using DESeq2, were clustered by K-means. Log-normalized expression values are shown, scaled by row. **(D)** Heatmap of differentially abundant proteins in blister fluid quantified via NULISA (p < 0.05, |log_2_FC| > 1, t-test). NULISA Protein Quantification (NPQ) represents log2-transformed normalized counts. Color was scaled by row. **(E)** Box plots showing *IFNA1:IFNA13* and *IFNB1* blister fluid protein levels (NPQ) across diseases and skins (lesional and non-lesional). T-test was used for pairwise comparisons: Not Significant (ns), \**P* < 0.05, ** *P* < 0.01, *** *P* < 0.001). **(F)** % of keratinocyte subtypes (of total keratinocytes) per disease; colors scaled by cell type (column). **(G)** Pseudo bulk expression heatmap of IFN-I–induced genes across keratinocyte subtypes. **(H)** UMAP embedding of reclustered lymphocytes. **(I)** % of lymphocyte subtypes (of total lymphocytes) per disease; colors scaled by cell type (column). **(J)** Pseudo bulk expression heatmap of cytotoxic effector and Type-I Interferon response genes. Th1 cells were nearly absent in HC and Pso.

Unsupervised clustering of 79,261 high-quality single cells identified four major cell types: keratinocytes (52,608), melanocytes (3,614), lymphocytes (14,256), and myeloid cells (8,783) (Fig. 1B), with defining markers consistent with prior analyses^4^ (Fig. S1A).

## RNA and protein profiling reveal disease-specific immune signatures

To uncover inflammatory pathways underlying each condition, we performed differential expression analyses at both the transcriptomic and proteomic levels. Pseudo-bulk aggregation of scRNA-seq data identified 3,176 genes differentially expressed across lesional and non-lesional skin compared to healthy controls (p < 0.01, |log FC| > 1) (Fig. 1C). Unsupervised clustering of the top differentially expressed genes revealed a shared type I interferon (IFN-I) signature in CLE and DM, characterized by elevated *MX1, OAS1, IFI27*, and *IFIT1* expression, which was present even in non-lesional skin (Fig. S1B).

Despite low transcript levels of IFN-I genes, corresponding proteins were robustly elevated in lesional CLE and DM skin, with modest increases also detected in non-lesional samples, suggesting a subclinical IFN-I milieu (Fig. 1D, E). IFN–inducible chemokines (*CXCL9, CXCL10, CXCL11*) were upregulated to varying degrees across all diseases, at both RNA (Fig. 1C) and protein levels (Fig. 1D). In contrast, psoriasis exhibited a dominant IL-17 axis, with marked upregulation of *CCL20, KRT6A, KRT16*, and *S100* family alarmins in lesional skin, some of which (e.g., *SPRR2A, S100A9, IL36G*) were also elevated in non-lesional samples (Fig. 1C (RNA) ,1D and S1C (Protein)).

## Basal keratinocytes in photosensitive skin exhibit a type I interferon signature

Keratinocytes clustered into six transcriptionally distinct subtypes, including basal, suprabasal spinous, granular, eccrine, and a heterogeneous group of keratinocytes at varying stages of differentiation, distinguished by high expression of *KRT6A*, *KRT6B*, *KRT16*, and *KRT17*. These genes are associated with wound healing and skin injury response, and hence we labeled them as “activated” as in previous studies^6^ (Fig. S1A). The proportions of these subtypes varied significantly across diseases (*χ*^2^ test p < 10^-15^) (Fig. 1F, Table S3), with activated keratinocytes enriched in psoriasis, CLE, and DM, consistent with heightened epithelial injury (Fig. S1D, scCODA FDR < 0.01). Granular keratinocytes were proportionally reduced in psoriasis, CLE, and vitiligo, whereas DM uniquely exhibited a pronounced loss of spinous keratinocytes, highlighting disease-specific alterations in keratinocyte differentiation and activation (Fig. 1F, Fig. S1D).

Strikingly, basal and suprabasal keratinocytes from lesional CLE and DM skin exhibited a transcriptional signature, marking a robust IFN-I response, absent in vitiligo and psoriasis (Fig. 1G). Notably, this signature was also detectable, albeit at lower levels, in non-lesional CLE and DM skin, suggesting a pre-existing IFN-I-primed state in keratinocytes from photosensitive diseases (Fig. 1G).

## Photosensitive diseases are enriched for cytotoxic CD4**^+^** T cells

Re-clustering of lymphocytes identified 12 transcriptionally distinct subsets, including Th1 CD4^+^ T cells, Th2-like naive CD4^+^ T cells (*IL7R, IL13, PTGDR2*), *FOXP3*^+^ regulatory T cells (Tregs), and three subsets of CD8^+^ T cells distinguished by *GZMB, GZMK*, or *IL-17* expression (Tc17), as well as mature and immature natural killer (NK) cells, γδ T cells, and an unlabeled low-transcript *CD3G*^+^ population (Fig. 1H, S1E).

Lymphocyte composition varied significantly across diseases (*χ*^2^ test p < 2.2 × 10^−1^ , Fig. 1I, Table S3). Tc17 cells were expanded in psoriasis while cytotoxic CD8 T cells (GZMB or GZMK high) were expanded in DM, CLE and vitiligo (Fig. 1I, S1F, scCODA FDR < 0.01). GZMK high CD8 T cells displayed intermediate GZMB expression but high expression of *GZMK, GZMA*, and *GZMH* (Figure S1E). A similar population of GZMK^+^ GZMB^+^ CD8^+^ T cells was recently described in synovial fluid and tissue of patients with rheumatoid arthritis^7^

Most notably, CLE and DM lesions were enriched for Th1-skewed CD4^+^ T cells that, unlike their counterparts in vitiligo, expressed cytotoxic effector molecules including *GZMB, GZMK*, and *PRF1*, a population nearly absent in healthy and psoriasis skin (Fig. 1I, 1J, S1F, scCODA FDR < 0.1). This was most prominent in DM, where Th1 cells had the highest expression of IFN-I response transcriptional signature (Fig. 1J).

In contrast, Th2-like naive CD4^+^ T cells and immature NK cells were depleted across all diseases relative to controls, reflecting a shift toward an activated, cytotoxic immune environment (Fig. 1I, S1F).

## Diverse myeloid cell states identified across inflammatory skin diseases

To define the heterogeneity of myeloid cells across inflammatory skin diseases, we re-clustered 8,783 myeloid cells into 8 transcriptionally distinct subpopulations (Fig. 2A). Two clusters expressed high levels of CD207 (Langerin), *CD1A*, and *HLA-DQB2*, and were annotated as Langerhans cells (LC). These differed in expression of *FCGBP* and *CSF1R.* Two clusters displayed strong expression of canonical conventional dendritic cell (DC) markers for cDC1 (*CLEC9A, XCR1,* and *CADM1) and cDC2 (SIRPA, FCE1RA, CLEC10A,* and *CD1C)* ^8^. One cluster had high levels of plasmacytoid DCs (pDCs) markers: *IL3RA (CD123), LILRA4 (ILT7),* and *IRF7* ^8, 9, 10^. Another cluster had high expression of monocytic lineage markers such as *MAFB, CD14, FCGR1A, and FCGR3A* (Fig. 2B). Given the combined DC and macrophage marker expression (*ITGAX*, *ITGAM*, *CD163*, *CD209*), we labeled this cluster CD14^+^ myeloid cells (CD14-MCs) to reflect its monocytic origin (Fig. S2A).

**Figure 2.**
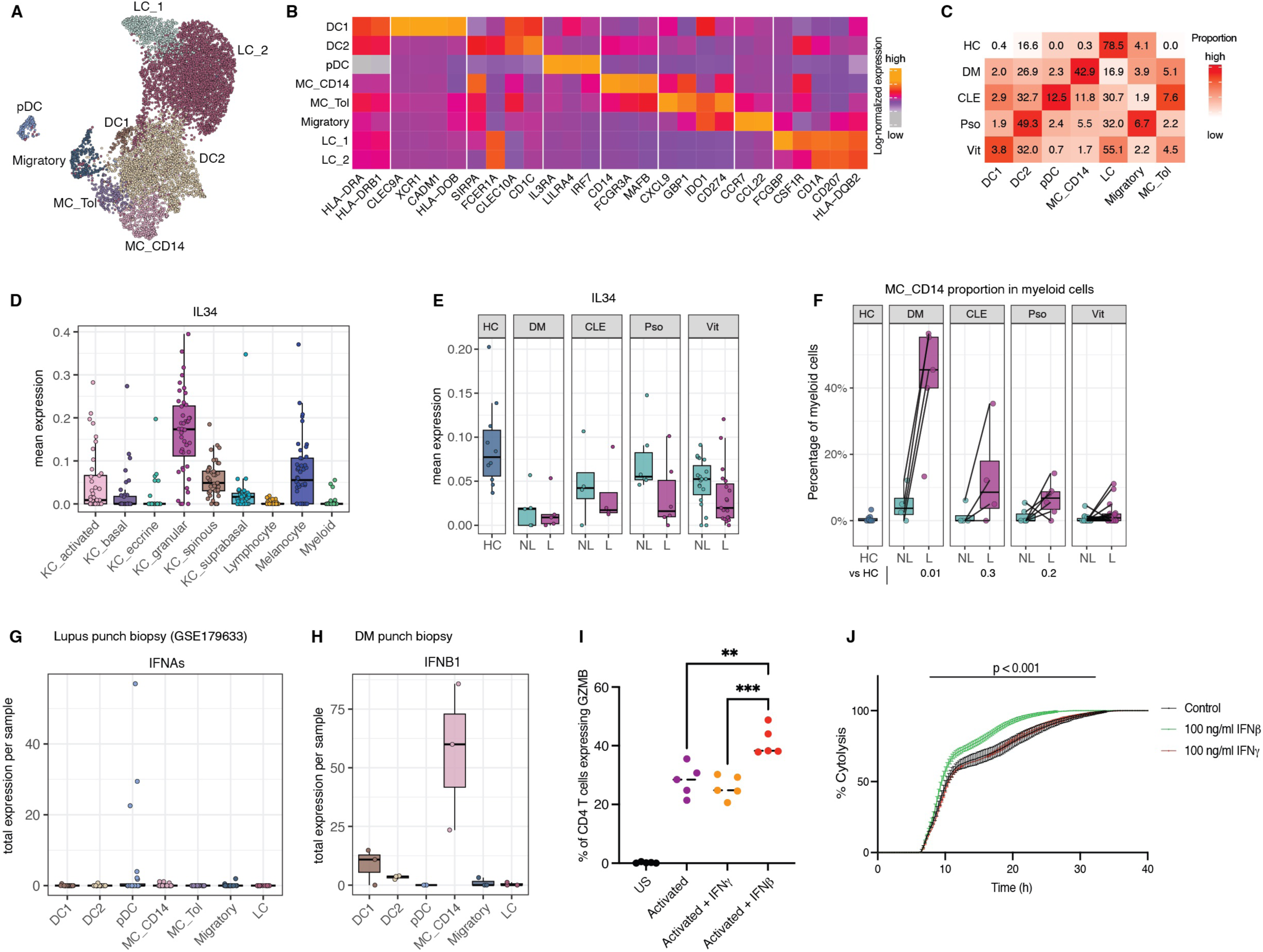
Disease-specific myeloid subsets and IFN-I–associated cytotoxicity in photosensitive skin disease. **(A)** UMAP embedding of reclustered myeloid cells. **(B)** Pseudo bulk expression heatmap of marker genes for myeloid subsets in scRNA-seq of blister biopsies. **(C)** Percentage of myeloid subtypes within all myeloid cells per disease; colors scaled by cell type (column). **(D,E)** Box plots showing IL34 pseudo bulk expression aggregated per cell type for each sample (D)**, or aggregated per sample (E);** each dot represents a sample. **(F)** Box plot showing percentages of CD14^+^ myeloid cells within total myeloid cells per sample, across skin conditions. Statistical significance was assessed using scCODA, with KC_suprabasal as the reference cell type. **(G, H)** Bar plots showing total log-normalized transcript counts for IFNAs (G) and IFNB1 (H) in myeloid cell subsets per sample in punch biopsies from public dataset GSE179633 (G), and three DM patients (H). Each dot represents a sample **(I)** Flow cytometry dot plot showing the frequency of anti-CD3/CD28 activated GZMB^+^ CD4^+^ T cells, treated with IFN-β or IFN-γ (n = 5 donors); Statistical analysis was performed using one-way ANOVA followed by Tukey’s multiple comparison test (** *P* < 0.01, *** *P* < 0.001). **(J)** Line plot of real-time cytolysis of solitomab treated BT20 cells co-cultured with CD4^+^ T cells pre-treated with IFN-β, IFN-γ, or control; data shown as mean ± SD; statistical comparison by one-way ANOVA followed by Tukey’s multiple comparison test (**p** < 0.001).

Finally, two migratory clusters were identified: one expressing *CCR7* and *CCL22*, consistent with lymph node-homing DCs^11^, and a myeloid population expression higher levels of tolerogenic genes (Myeloid-Tol): *CXCL9*, *GBP1*, *GBP5*, *IDO1*, *CD274* (PDL1), and *NECTIN2*. A similar cell population previously shown to have immunomodulation function^12^ (Fig. 2B).

## Myeloid cells from diseased skin display shared and disease-specific changes compared with healthy skin

Myeloid cell proportions were significantly altered across all four inflammatory skin diseases (p < 2.2×10^-16^, *χ*^2^ test, Fig. 2C, Table S3). Myeloid-Tol cells, rarely observed in healthy skin, showed consistent expansion in diseased samples (FDR < 0.1, scCODA, Fig. S2B). Conversely, we observed a consistent reduction in LCs compared to healthy skin (FDR < 0.2, scCODA, Fig. S2B).

LC depletion aligns with prior studies indicating that LCs migrate to skin-draining lymph nodes upon activation following tissue injury^13, 14^. However, the underlying mechanisms regulating this process remain poorly understood. Prior studies have established IL-34 as a critical factor not only for the development, but also for the differentiation and maintenance of tissue resident LCs, including those in the skin^15, 16^. Our scRNA-seq data revealed that IL34 is predominantly expressed by granular keratinocytes, with lower expression spinous keratinocytes and differentiated melanocytes (Fig. 2D). Notably, IL34 RNA levels were consistently downregulated in lesional skin across all four diseases (p < 0.001; Fig. 2E). This reduction reflects both the loss of IL34-expressing keratinocyte and melanocyte subsets and diminished per-cell IL34 transcript levels. The resulting decrease in local IL34 levels may impair LC survival, contributing to their depletion in lesional skin.

In contrast to LC loss, DC2 subsets were expanded in disease, especially in psoriasis, where they accounted for up to 50% of myeloid cells in lesional skin (Fig. S2B, Table S3). This finding is consistent with previous studies demonstrating DC2 accumulation in human psoriasis and mouse models, where DC2 depletion reduces IL-17–mediated psoriasiform inflammation^17^.

DC1 cells, known for their capacity to cross-present antigens and prime CD8+ T cells^18^ – were specifically expanded in vitiligo, where they reached 3.8% of lesional myeloid cells, compared with 0.4% in healthy skin (FDR < 0.3, scCODA, Fig. 2C, S2B). This expansion is consistent with our prior work indicating that DC1 depletion protects against disease in a vaccinia virus-induced mouse model of vitiligo^19^.

## Expansion of type I interferon–producing myeloid cells distinguishes photosensitive from photoresponsive skin disease

A key distinction revealed by our comparative single-cell analysis of inflammatory skin diseases was the expansion of type I interferon–producing myeloid subsets specifically in photosensitive dermatoses. Two such populations stood out: plasmacytoid dendritic cells (pDCs) in CLE and CD14^+^ myeloid cells (MC_CD14) in DM (Fig. 2C, Table S3). pDCs were markedly enriched in lesional CLE, comprising 12.5% of the total myeloid compartment, compared to less than 2.5% in DM and psoriasis, and 0.7% in vitiligo, while they were virtually undetectable in healthy skin. Similarly, CD14^+^ MC cells, scarce in healthy skin (<0.3%), represented 43% of myeloid cells in DM and 12% in CLE lesions (Fig. 2F, S2B, scCODA FDR < 0.01, Table S3).

The role of pDCs in CLE pathogenesis is well established, largely attributed to their secretion of IFN-⍰^20^. Notably, a recent phase 2 randomized controlled trial in CLE demonstrated clinical benefit from targeting pDCs via a monoclonal antibody against CLEC4C (BDCA2), a pDC-specific surface marker^21^. Although *IFNA* transcripts were not detected in our scRNA-seq dataset—likely due to technical dropout—a deeply sequenced public dataset (GSE179633^22^, methods) from full-thickness CLE skin confirmed pDCs as the predominant source of *IFNA* (Fig. 2G, Fig. S2C).

CD14-MC cells, by contrast, emerged as a prominent source of IFNB1 in DM. While single-cell detection of *IFNB1* is limited, we identified *IFNB1* transcripts primarily within the CD14^+^ cluster in our dataset (Fig. S2D). We validated this finding in a second independent scRNA-seq dataset from full-thickness skin biopsies of lesional skin of three additional DM patients (Methods, Fig. 2H^23^), and *IFNB1* was again predominantly expressed by CD14^+^ cells (Fig. S2E,F).

Together, these findings identify a disease-specific expansion of type I interferon–producing myeloid subsets in photosensitive dermatoses: pDCs in CLE producing IFN-α and CD14^+^ MC cells in DM, and to a lesser extent in lupus, producing IFN**-**β. Consistent with their transcriptional profiles and relative abundance, protein levels of IFN-α and IFN-β were significantly elevated in interstitial fluid from CLE and DM lesional skin, respectively (Fig. 1E, Fig. S1A), reinforcing their central role in driving photosensitivity-associated inflammation.

## CD4 T cells exhibit enhanced cytotoxicity in response to type I, but not type II, interferons

Th1 cells, defined by high expression of *TBX21*, *IFNG*, and *CXCR3* (Fig. S1E), were markedly enriched in photosensitive lesions, with the greatest expansion observed in DM. In lesional DM skin, these Th1 cells also expressed the highest levels of cytotoxic markers, most notably *PRF1* and *GZMB* (Fig. 1I,J). Similar cytotoxic CD4^+^ T-cell subsets have been reported to mediate antiviral activity through granzyme- and perforin-dependent killing ^24^.

The predominance of cytotoxic Th1 cells in DM prompted us to ask whether type I interferon augments their cytolytic potential. CD4^+^ T cells from healthy donors were activated with anti-CD3/CD28 for 72 h and then cultured with either IFN-β or IFN-*γ*. Activation alone significantly increased *GZMB* expression (p *<* 0.001); while addition of IFN-β, but not IFN-*γ*, further enhanced *GZMB* levels (*p* < 0.01; Fig. 2I).

To determine whether this translated into greater killing capacity, we evaluated cytotoxicity using a lymphocyte cytotoxicity assay^25^ (Agilent Real-Time Immune Cell Killing Assay). Primary CD4^+^ T cells from healthy donors were incubated with solitomab (CD3-EpCAM), followed by incubation with BT20, an epithelial ductal carcinoma cell line (ATCC). This approach forces CD4^+^ T and their BT20 targets to interact via the bispecific antibody, permitting isolated measurement of cytotoxic capacity. Consistent with the increase in GZMB expression, pre-treatment with IFN-β, but not IFN-*γ*, significantly enhanced the cytotoxic function of CD4^+^ T cells (Fig. 2J). Collectively, these results indicate that type I interferon selectively enhances the cytotoxic program and effector function of Th1 CD4^+^ T cells, a mechanism that likely contributes to tissue damage in photosensitive skin disease.

## Spatial transcriptomics of autoimmune skin lesions

To better understand the spatial context and interactions of the two cell types most characteristic of photosensitive disease lesions, CD14^+^ myeloid cells and cytotoxic CD4^+^ T cells, we turned to spatial transcriptomics using Sequential fluorescence in situ hybridization (seqFISH), which enable the simultaneous detection of hundreds of gene transcripts with subcellular resolution^3^.

We designed a panel of 511 genes informed by our single-cell RNA-seq analysis. This panel included: (i) canonical markers defining the clusters identified in scRNA-seq; (ii) established cell-type markers from prior studies; (iii) 60 cytokines and their receptors to enable inference of cell-cell communication; (iv) differentially expressed genes from prior studies^4^ and (v) housekeeping genes as internal controls. In addition, we included cytokines and dysregulated transcripts identified in our own differential expression analyses.

For seqFISH analysis, we obtained 4 mm punch skin biopsies from an additional cohort of seven patients with active disease (2 DM, 2 CLE, 2 psoriasis, and 1 vitiligo), yielding 13 spatial transcriptomics datasets (two tissue sections per biopsy, except for one case). After segmentation and assigning transcripts to cells, we detected a total of 155,440 cells and an average of 594 transcripts per cell across our samples (Methods).

Unsupervised clustering of seqFISH data recapitulated all major cell types previously defined in our scRNA-seq dataset and additionally resolved several dermal populations not captured in suction blister samples (Methods, Fig. 3A). The spatial distribution of these clusters across samples closely matched known skin architecture, confirming accurate mapping of cellular identities (Fig. 3B, Supplementary Material 1).

**Figure 3.**
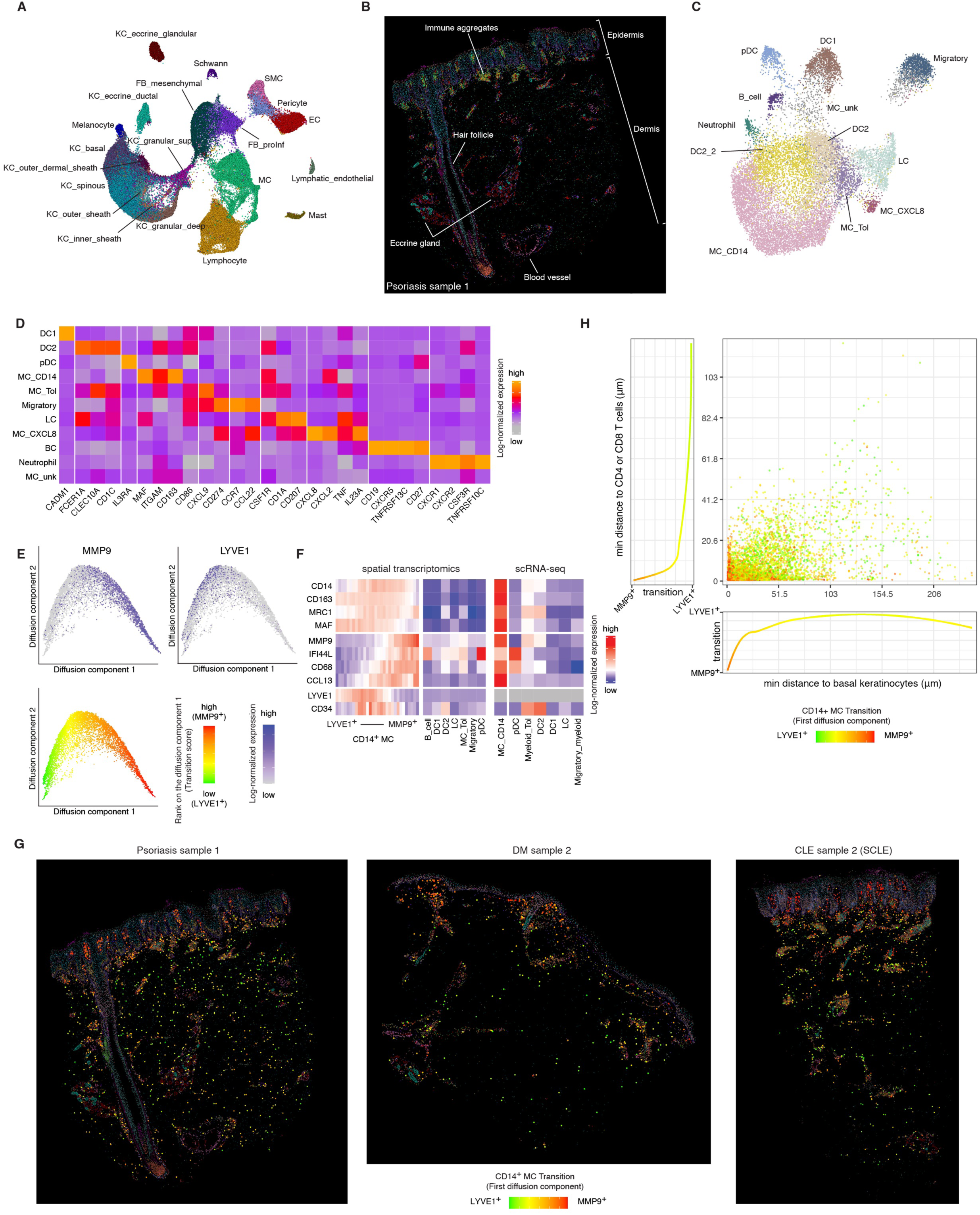
Spatial transcriptomics reveals spatial stratification and transcriptional heterogeneity of CD14^+^ myeloid cells in autoimmune skin lesions. **(A)** UMAP embedding of 155,440 cells profiled by seqFISH across 13 skin sections. **(B)** Spatial map showing key skin features captured in a representative psoriasis tissue section; each dot represents a cell; colored by cell type **(C)** UMAP embedding of reclustered seqFISH myeloid lineage. **(D)** Pseudo bulk expression heatmap of marker genes for myeloid subclusters in seqFISH of punch biopsies. **(E)** Diffusion map of CD14^+^ myeloid cells identify a transcriptional continuum from LYVE1^+^MMP9⁻ to MMP9^+^LYVE1⁻ states. Color intensity indicates log-normalized single cell expression (Upper two panels) or the first diffusion component score (bottom panel). **(F)** Heatmap of selected marker genes in myeloid cell subtypes. seqFISH (left and middle). Expression of CD14+ cells ordered by their first diffusion component and aggregated into 50 equal-sized bins for the mean expression level of each bin (left). Right plot shows expression in myeloid subtypes in the blister fluid scRNA-seq data. Left and middle were scaled together. **(G)** Spatial embeddings of seqFISH data showing CD14+ myeloid cells colored by their first diffusion component in lesional tissue sections from psoriasis, DM and CLE. **(H)** Scatter plot showing minimum distance in μm from a CD14+ cell to a basal keratinocyte (x-axis) or a CD8/CD4 T cell (y-axis) colored by their projection onto the first diffusion component; marginal plots show the smoothed values of the first diffusion component of CD14+ cells projected onto the x-axis (bottom) or the y-axis (left).

To resolve finer cellular subtypes, we performed secondary clustering within each major lineage (Fig. 3C, S3A-D, Supplementary Material 2). Reclustering of the myeloid compartment identified populations corresponding to DC1, DC2, CD14^+^ Myeloid cells (MC_CD14) tolerogenic myeloid cells (Myeloid-Tol), Langerhans cells (LC), and plasmacytoid dendritic cells (pDCs), consistent with our scRNA-seq findings. Notably, two small additional populations emerged that were not detected in blister biopsy datasets: a neutrophil cluster (263 cells) characterized by expression of *CSF3R, CXCR1, CXCR2,* and *TNFRSF10C (TRAILR3, DCR1)* (Fig. 3D); and an epidermal LC-like subset (242 cells, labeled MC_CXCL8), primarily derived from samples with hyperproliferative keratinocyte, expressing canonical LC markers (*CD1A*, *CD207)*, along with proinflammatory cytokines *CXCL8*, *CXCL2* and *IL23A* (Fig. 3D, S3E).

In addition to immune populations, seqFISH enabled characterization of cell subsets not captured in suction blister samples. We identified five fibroblast subtypes, including mesenchymal fibroblasts, which were evenly distributed throughout the reticular dermis. Notably, two distinct pro-inflammatory fibroblast subsets, characterized by expression of *CXCL2*, *CXCL3* and *CCL19*^26^, localized to separate dermal compartments. Superficial pro-inflammatory fibroblasts (FB_proInf_sup) resided just beneath the dermal-epidermal junction, whereas deep pro-inflammatory fibroblasts (FB_proInf_deep) were enriched within immune aggregates and surrounding eccrine structures (Fig S3F). These subsets were primarily distinguished by their chemokine expression profiles, with FB_proInf_sup expressing higher levels of *CCL8*, *CCL2*, and *CXCL2*, and FB_proInf_deep was enriched for *CCL19* and *CXCL12* (Fig S3B).

## Spatial transcriptomics reveals spatial stratification and inflammatory heterogeneity of CD14+ myeloid cells

To investigate the spatial organization and inflammatory states of CD14^+^ myeloid cells in photosensitive skin lesions, we focused our analysis on this population. Marker gene analysis revealed the expression of *LYVE1* and *CD34*, which were notably absent in CD14^+^ cells captured via the suction blistering method. This suggested the presence of previously undetected CD14^+^ subsets that may be excluded from blister biopsies.

CD14^+^ MCs appeared to form a transcriptional continuum, so we embedded CD14+ cells using a diffusion map (Methods; Fig. 3E). While all cells shared a core transcriptional profile marked by high expression levels of *CD14*, *CD163*, *MRC1*, and *MAF*, key differences emerged through the trajectory. Specifically, cells at one end of the trajectory expressed low levels of *MMP9*, *CD68*, and *IFI44L*, genes we had found highly expressed in CD14^+^ cells isolated via suction blistering. These cells expressed high levels of *LYVE1* and *CD34*, which were virtually absent in previously captured CD14^+^ cells (Fig. 3F). This suggested that certain CD14^+^ cells were not captured by blistering. On the other hand, cells at the opposite end of the trajectory expressed high levels of *MMP9*, *CD68*, and *IFI44L* that more closely resembled CD14^+^ cells observed in suction blisters.

CD14^+^ cells along the transition localized at different domains in the skin. While LYVE1^+^CD14^+^ cells were distributed deeper within the dermis, CD14^+^MMP9^+^ cells were generally more superficial and near the dermal-epidermal junction and immune aggregates (Fig. 3G,H, S4A). This data suggested a state transition in which cells acquire a proinflammatory profile by upregulating *MMP9* and downregulating *LYVE1* (Fig. S4B).

A key factor influencing the selective capture of MMP9^+^CD14^+^ cells in blister biopsies appears to be MMP9’s function, which encodes a matrix metallopeptidase that facilitates extracellular matrix degradation and immune cell migration. CD14^+^ cells with high MMP9 and low LYVE1 expression were likely to be more easily detached from skin and thus more likely to be sampled by suction blistering. In contrast, CD14^+^ cells with high expression of LYVE1 likely remain anchored within the dermis and are not efficiently captured by the blister biopsy technique, because LYVE1 encodes a protein that binds hyaluronic acid, which is a major component of the dermal extracellular matrix.

## Colocalization analysis reveals spatial proximity of inflammatory CD14^+^ myeloid cells and cytotoxic CD4^+^T cells in photosensitive skin lesions

To quantitatively assess spatial relationships between CD14^+^ myeloid cells and other populations, particularly the expanded Th1 cytotoxic CD4^+^ T cell population observed in DM lesions (Fig. S4C,D), we developed a spatial metric termed colocalization strength. This metric integrates both the density and proximity of one cell type relative to a reference cell type (Methods). Our approach models each cell from the reference type (e.g. CD4^+^ T cells) as a Gaussian distribution centered at its centroid. For every cell of interest, we compute a co-localization score by summing Gaussian-weighted distances to all reference-type cells within a defined radius. Changing the radius enables capturing colocalization at varying spatial scales (Methods, Fig. S4E). Averaging these scores across all cells of a given type provides a measure of colocalization between two populations (Methods). Notably, this metric is direction dependent, reflecting asymmetric spatial distributions of interacting cell populations.

We first validated our method by confirming known anatomical colocalization pattern in skin. For example, we observed expected clustering of glandular and ductal eccrine keratinocytes, Schwann cells, and pericytes within eccrine gland regions (box 1, Fig. 4A). Similarly, clustering accurately grouped the known distinct layers of the hair follicle sheath (box 2, Fig. 4A). Additionally, epidermal clusters accurately grouped keratinocytes from different epidermal layers, melanocytes, Langerhans cells, and Tc17 cells^27^ (box 3, Fig. 4A). We also observed endothelial cells and fibroblasts bridging dermal structures and immune cell clusters, while proinflammatory *CCL19*^+^ fibroblasts were concentrated in immune rich regions distinguishing them from dermal mesenchymal fibroblasts (box 4 and 5, Fig. 4A).

**Figure 4.**
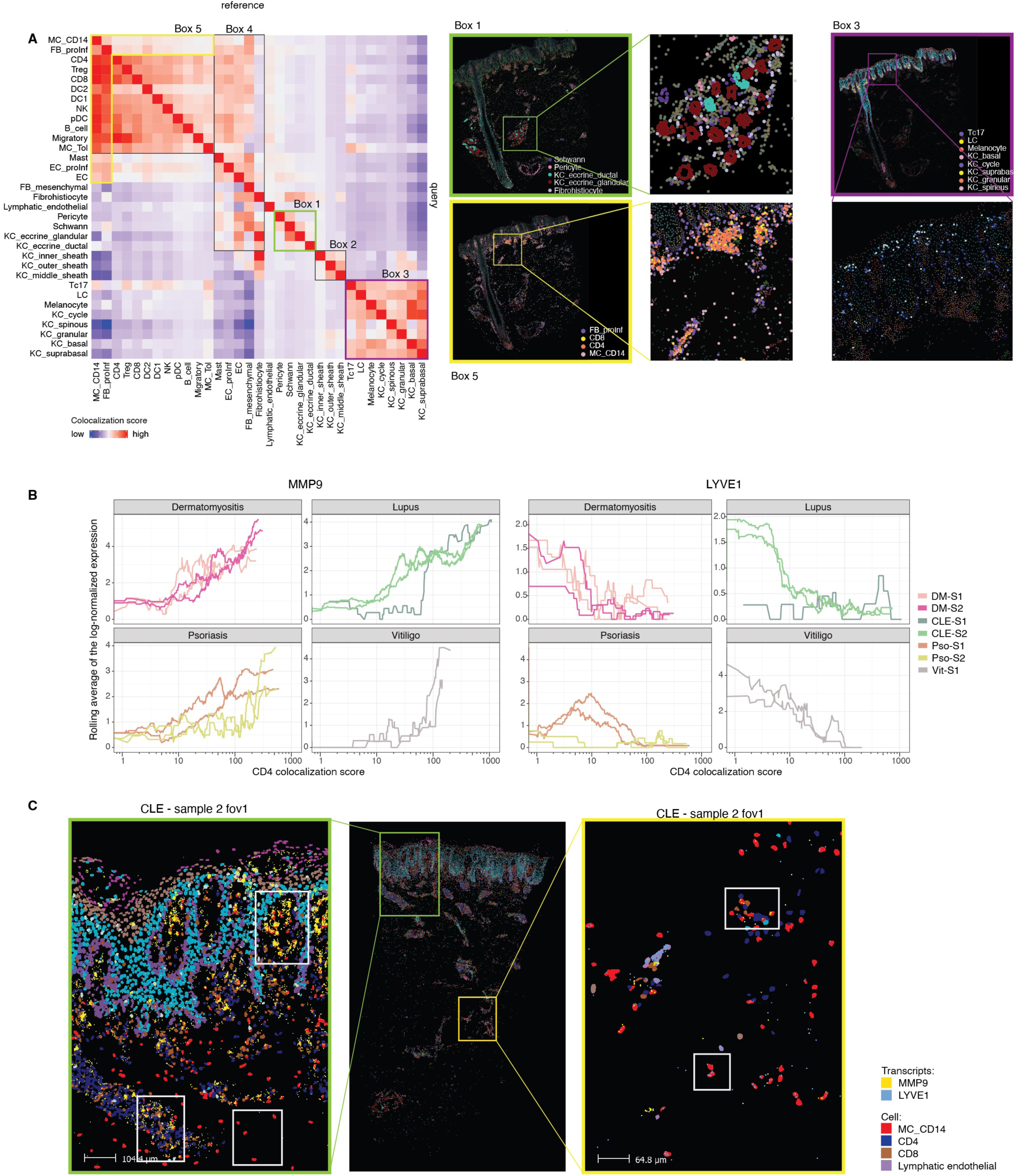
Spatial colocalization analysis reveals activation states of CD14^+^ myeloid cells associated with proximity to CD4^+^ T cells in autoimmune lesions. **(A)** Left: Heatmap of cell-type colocalization scores across all spatial transcriptomics slides, calculated using a Gaussian-weighted distance model (see Methods). Score shows the degree of query cell type (row) locating within the region of the reference cell type (column). Boxes 1–5 highlight known anatomical relationships: (1) Eccrine gland: glandular and ductal keratinocytes with Schwann cells and pericytes; (2) Hair follicle: concentric layers of hair follicle sheath keratinocytes; (3) Epidermis: epidermal keratinocytes with melanocytes, Langerhans cells, and Tc17 cells; (4) Dermis: mesenchymal fibroblasts with dermal structures; (5) Immune aggregates: proinflammatory fibroblasts with immune cells. Right: Zoom-ins in representative tissue sections of the highlighted clusters. **(B)** Gene in CD14^+^ myeloid cells expression changes as a function of proximity (colocalization score) to CD4^+^ T cells aggregated for each slide; smoothed with rolling average. **(C)** High-resolution image of CLE sample 2 showing zoom-ins of superficial and deep immune aggregates. Key cell types are displayed, with overlaid transcripts for LYVE1 (blue) and MMP9 (yellow). Lymphatic endothelial cells, typically associated with LYVE1 expression, are shown for spatial reference.

Subsequent analysis uncovered dynamic gene expression changes in CD14^+^ myeloid cells associated with proximity to CD4^+^ T cells. Specifically, CD14^+^ cells displayed increased *MMP9* and decreased *LYVE1* expression as they approached CD4^+^ T cells (Fig. 4B,C, S4F). Elevated *MMP9* suggests enhanced extracellular matrix remodeling capacity, supporting cell migration and facilitating immune synapse formation with CD4^+^ T cells^28, 29^. In contrast, reduced *LYVE1* expression reflects a loss of tissue-resident homeostasis and a shift toward a more migratory, inflammatory phenotype^30^, consistent with activation in type I interferon–rich microenvironments.

## CD14^+^ myeloid cells polarize from M2 to M1 as they migrate toward the epidermis in photosensitive skin diseases

To further dissect the transcriptional heterogeneity of CD14^+^ myeloid cells, beyond the limitations of our seqFISH panel which had a restricted by probe set and the suction blister biopsies, which underrepresent deep dermal infiltrates, we reanalyzed a previously published (GSE179633), deeply sequenced single-cell RNA-seq dataset generated from separately isolated dermal and epidermal compartments of skin biopsies from healthy controls and lesional CLE samples from patients with or without systemic lupus erythematosus^22^. After Harmony-based batch correction (Methods), we focused on refining the myeloid and lymphocyte compartments to enable high-resolution characterization.

As expected, pDCs were markedly enriched in both discoid lupus erythematosus (DLE) and SLE relative to healthy skin. Importantly, CD14^+^ cells demonstrated significant expansion in the epidermal samples but not in dermal samples, consistent with our spatial data (Table S3; Fig. 5A). Targeted clustering of CD14^+^ cells revealed a similar continuous transcriptional trajectory to what we had observed above, with *MMP9* and *LYVE1* defining opposite poles along the first diffusion map dimension (Fig. 5B).

**Figure 5.**
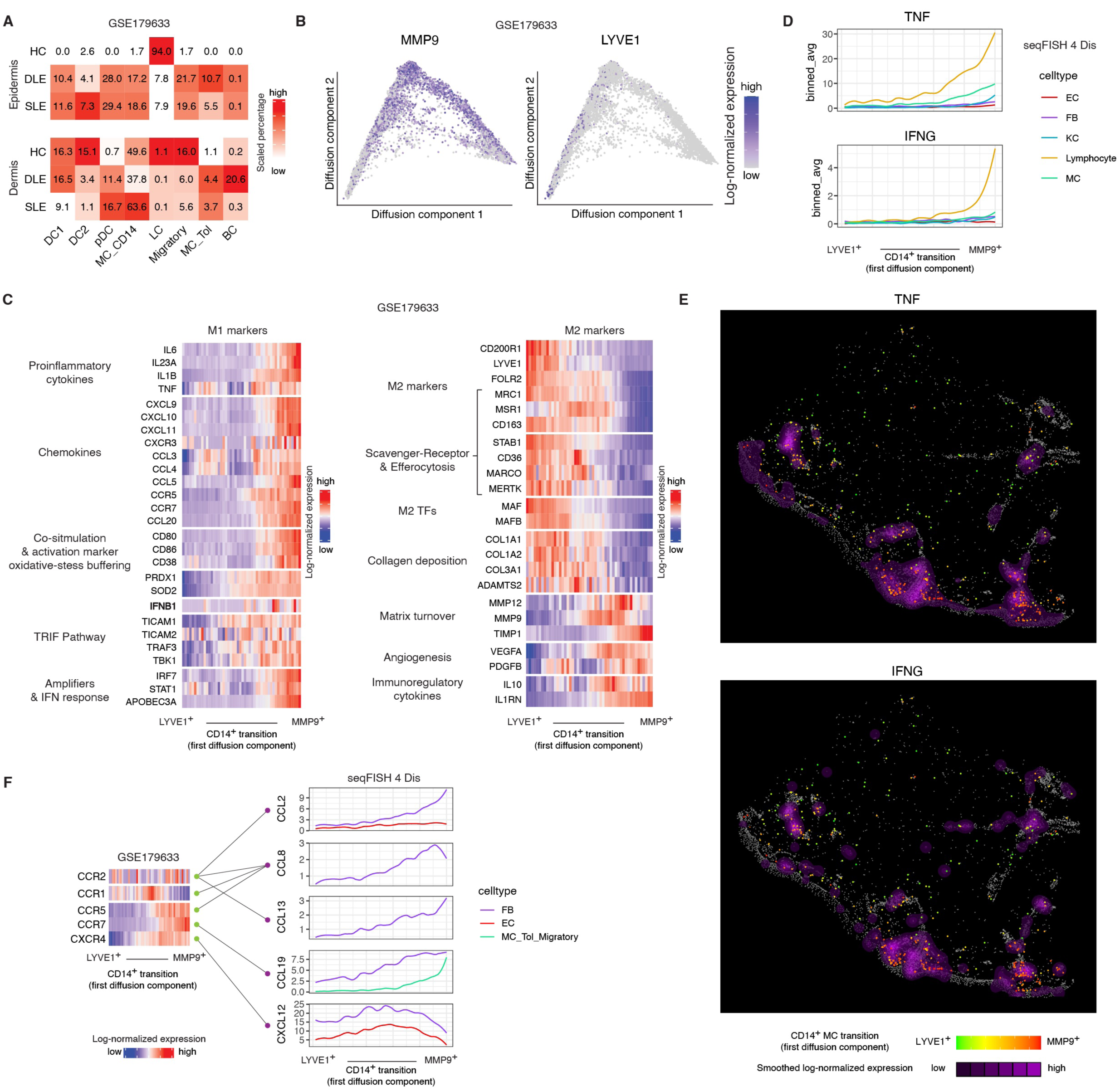
CD14^+^ cells transition from an M2-like to an M1-like state and their skin localization is established by a multi-chemokine fibroblast gradient. **(A)** Percentage of myeloid cells subtypes within all myeloid cells per condition in epidermal (top) and dermal (bottom) punch biopsy samples from reanalyzed data in GSE179633; colors scaled by cell type (column). **(B)** Log-normalized expression of MMP9 (left) and LYVE1 (right) expression in CD14^+^ cells embedded in a diffusion map. **(C)** Heatmap showing M1 (left) and M2 (right) markers expression in CD14^+^ cells ordered by the first diffusion component (transition) and aggregated into 50 bins (144 cells per bin). **(D)** Local expression of TNF (top) and IFNγ (bottom) by cells within 400px (∼41μm) of CD14^+^ cells. CD14^+^ cells were ordered by their first diffusion component (transition), and averaged in bins of 193 CD14^+^ cells. **(E)** Expression of TNF (top) and IFNγ (bottom) in a DM lesional sample, smoothed with gaussian-kernel sd=400px (∼41μm); CD14^+^ cells are overlayed on top and colored according to their projection onto the first diffusion component (transition). **(F)** Left: Expression of detected chemokine receptors in CD14^+^ cells from the GSE179633 dataset, shown along the first diffusion component. CD14^+^ cells grouped into 50 bins (144 cells per bin) as in (B). Right: seqFISH expression of corresponding ligands in different cell types within 400px (∼41μm) of CD14^+^ cells, averaged per bin along the same diffusion component ordering as in (D).

Differential gene expression analysis across the continuum identified 3,234 significantly differentially expressed genes (padj < 0.05 & |avg_log2FC| > 1; Fig. S5A). CD14^+^ cells at the *MMP9+* end exhibited a transcriptional program consistent with M1 polarization, including elevated expression of proinflammatory cytokines (*IL6*, *IL1B*, *TNF*), whereas cells at the *LYVE*^+^ end displayed an M2-like signature (p < 0.01) (Fig. 5C).

Since moDC/Macs are highly plastic and can switch between M1- and M2-like states in response to changes in the local environment, we explored the spatial pattern of M1 and M2 inducers relative to the location of MMP9^+^CD14^+^ cells and LYVE1^+^CD14^+^ cells. M2-inducer TGFB1 was expressed constitutively by fibroblasts. Another M2-inducer, glucocorticoid that is produced by the adrenal cortex, is always present in skin, shaping an M2 default state. In contrast, M1 inducers, IFNG and TNF, were broadly produced by immune cells. Spatially, MMP9^+^CD14^+^ cells colocalized with high TNF and IFNG production at the dermal-epidermal junction and immune aggregates (Fig. 5D, E), likely tilting the balance of CD14^+^ cells toward an M1 phenotype.

To identify mechanisms guiding the spatial positioning of CD14^+^ cells along their transcriptional transition, we analyzed chemokine receptor expression in the public dataset. CD14^+^ cells expressed five chemokine receptors: CCR1, CCR2, CCR5, CCR7 and CXCR4. CCR1 and CXCR4 were expressed midpoint in the transition, while CCR5 and CCR7 increased towards the MMP9^+^ end of the transition. Notably, CCR2, a known monocytic chemoattractant receptor^31^, was expressed at relatively low levels, with modest upregulation towards the MMP9^+^ end of the transition (Fig 5F left).

We next looked for cell types expressing ligands for these receptors. We find that fibroblasts, lymphocytes, myeloid cells (including CD14^+^ cells), and endothelial cells demonstrated high expression of these ligands (Figure S5B). To spatially localize ligand-producing cells near CD14^+^ cells, we leveraged our seqFISH dataset, which included a subset of these ligands (Fig. 5F).

Distinct expression patterns emerged among fibroblasts located within 400px (∼40μm) of CD14^+^ cells along their transition. CXCL12 a ligand for CXCR4, was predominately expressed by fibroblasts near CD14^+^ cells at the LYVE1 end of the trajectory. Notably, CXCR4 expression in CD14^+^ cells was also elevated closest to the LYVE1 compared to other receptors. In contrast, fibroblasts expressed CCL8 (CCR1, CCR2, CCR5 ligand) adjacent to CD14^+^ cells near the MMP9^+^ endpoint. CCL19 (CCR7 ligand) expression peaked at an intermediate stage while CCL2 (CCR2 ligand), reached highest expression towards the MMP9^+^ stage. These results suggests that pro-inflammatory fibroblasts in lesional skin orchestrate the spatial progression of CD14^+^ cells as they transition to an (MMP9^+^) inflammatory state through specific chemokine gradients (Figure 5F).

Endothelial cells adjacent to CD14^+^ cells expressed high levels of CXCL12 and CCL19 towards the midpoint of the transition (Fig. 5F).

Since many important CCR1 and CCR5 ligands were absent from our seqFISH panel, we examined their expression using again the GSE179633 dataset. Endothelial cells, particularly venous endothelial cells, highly expressed CCL14, CCL15 and CCL16, suggesting CCR1-mediated recruitment of CD14^+^ cells into skin, potentially differentiating into LYVE1^+^ or MMP9^+^ cells depending on inflammatory context (Fig. S5B). Given our steady-state analysis this alternative state transition progression to our proposed LYVE1^+^ to MMP9^+^ transition cannot be ruled out

T cells emerged as the major producers of CCL5, a key CCR5 ligand, pointing to their role in the formation of the observed CD14^+^MMP9^+^/T cell immune aggregates characteristic of CLE and DM lesions. Finally, CD14^+^ MCs expressed CCL2/3/4/7/8/13, suggesting self-recruitment through CCR1/2/5 (Fig. S5B).

## Type I interferon primes UVB-induced recruitment of MMP9^+^ CD14**^+^** myeloid cells in photosensitive skin and is abrogated by IFNAR blockade

Spatial and transcriptional profiling revealed a striking transition in CD14^+^ myeloid cells within inflamed skin, from tissue-resident, M2-like cells in the deep dermis to proinflammatory, M1-like cells enriched near the dermal–epidermal junction and immune aggregates. This phenotypic shift, marked by increased expression of *MMP9*, *IL1B*, and *TNF* alongside reduced *LYVE1* (Figure 5C), likely results in the expansion of CD14^+^MMP9^+^ cells observed in blister fluid samples of the two prototypical photosensitive diseases, CLE and DM, both of which are characterized by robust type I interferon (IFN-I) signatures (Fig. 1C).

Given that IFN-I signatures were also elevated in non-lesional skin from patients with CLE and DM, we hypothesized that type I interferon signaling may prime clinically uninvolved skin for an exaggerated immune response to UVB radiation. To test this, we conducted in vivo UVB photoprovocation in three healthy individuals and three patients with DM (Methods). Briefly, the minimal erythema dose (MED) was determined for each subject, after which clinically normal appearing, photoprotected skin on the lower back was irradiated with 1.5× MED of broadband UVB. Twenty-four hours post-irradiation, suction blister biopsies were obtained from both the UVB-treated and site-matched untreated skin for flow cytometric analysis (Fig. 6A). In healthy controls, CD14^+^ cells made up less than 10% of CD45^+^ immune cells in both irradiated and unirradiated skin. In contrast, UVB exposure in non-lesional DM skin triggered a robust infiltration of CD14^+^ cells, which comprised approximately 20% of total CD45^+^ immune cells (p < 0.001; Fig 6B, S6A).

**Figure 6.**
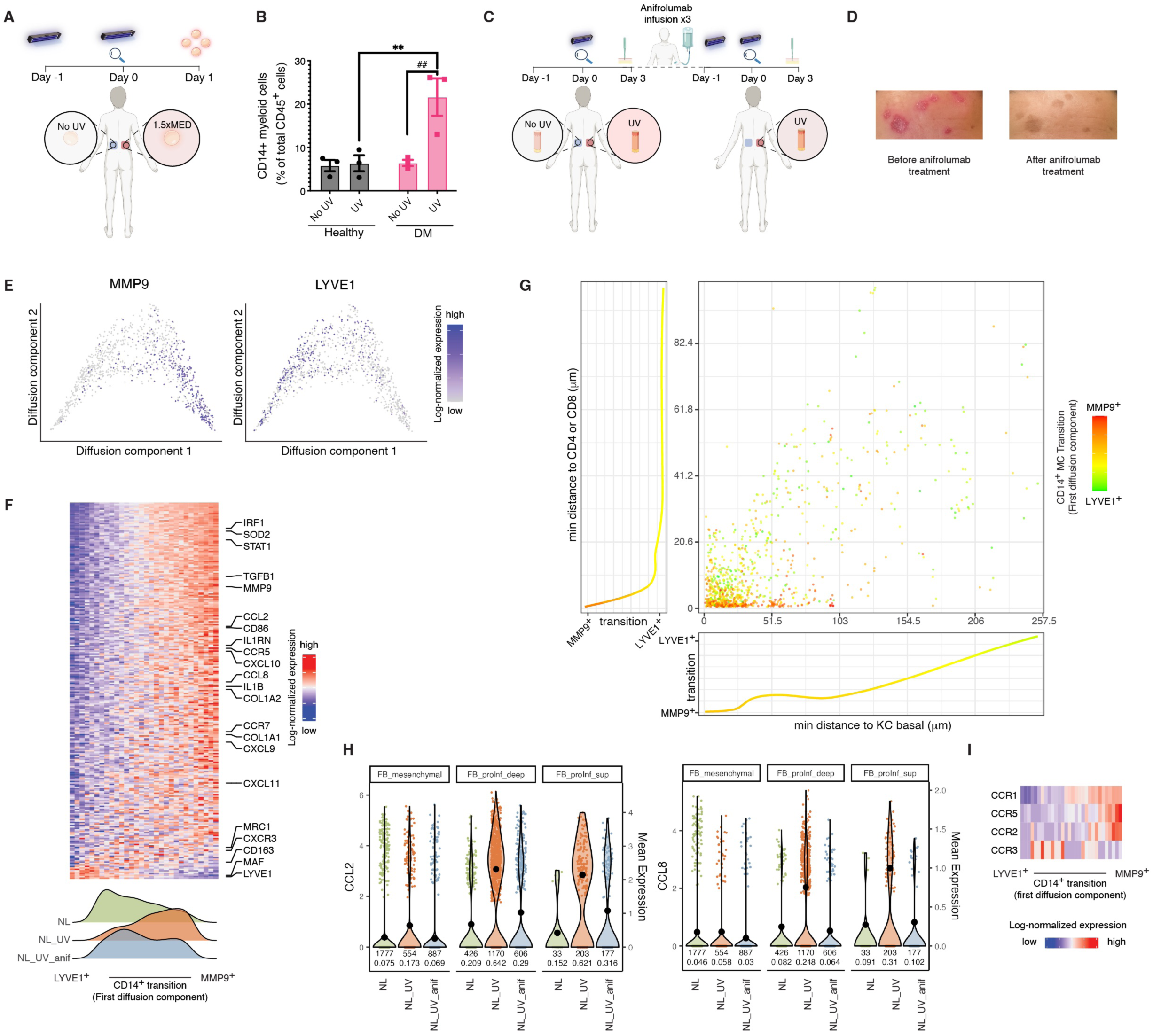
Type-I interferon signaling drives UVB-induced myeloid inflammation in DM, which is attenuated by therapeutic IFNAR blockade. **(A)** Schematic of the UVB photoprovocation protocol. Minimal erythema dose (MED) was determined for each subject, followed by exposure of non-lesional, photoprotected lower back skin to 1.5× MED UVB. Suction blister biopsies were collected 24 hours later from both irradiated and unirradiated sites. **(B)** Flow cytometric analysis of the percentage of CD14^+^ myeloid cells within CD45^+^ immune cells. p-values were computed using two-way ANOVA followed by Bonferroni-corrected pairwise t-tests (n = 3**). (C)** Schematic of anifrolumab intervention experiment. A patient with cutaneous lupus erythematosus (CLE) underwent UVB provocation before and after three monthly IV infusions of anifrolumab (anti-IFNAR1 monoclonal antibody). **(D)** Clinical images before and after anifrolumab treatment. CLASI activity score from 6 (before) to 0 (after). **(E)** Diffusion map of CD14^+^ cells obtained from seqFISH data of UVB-exposed skin. **(F)** Heatmap of differentially expressed genes (p < 0.01, tradeSeq) in CD14^+^ myeloid cells ordered by their first diffusion map component and their expression summed in 50-bin cells isolated from untreated (NL), UVB-treated (NL_UV), and UVB-treated post-anifrolumab (NL_UV_anf) skin. **(G)** Scatter plot showing minimum distance in μm from a CD14^+^ cell to a basal keratinocyte (x-axis) or a CD8/CD4 T cell (y-axis) colored by their projection onto the first diffusion component; marginal plots show mean of the first diffusion component of CD14^+^ cells projected onto the x-axis (bottom) or the y-axis (left) **(H)** Violin plots showing normalized expression of *CCL2* and *CCL8* in fibroblast subtypes. **(I)** Heatmap showing expression of chemokine receptors *CCR1*, *CCR2*, and *CCR5* in CD14^+^ myeloid cells ordered by the first diffusion component; each tile represents aggregated expression of 33 cells. (ns: not significant, * or # or ^ p < 0.05; ** or ## or ^^ p < 0.01, *** or ### or ^^^ p < 0.001, **** or #### or ^^^^ p < 0.0001).

To further validate the role of IFN-I signaling in this process, we leveraged an in vivo therapeutic intervention in a patient with active recalcitrant CLE who underwent UVB photoprovocation before and after three monthly infusions of anifrolumab, a monoclonal antibody targeting the type-I IFN receptor (IFNAR1) (Fig. 6C). Treatment with anifrolumab led to a marked increase in the patient’s MED (from 75 to 100 mJ/cm^2^; data not shown), as well as a complete resolution of clinical disease activity as reflected by a drop in the CLASI activity score (from 6 to 0; Fig. 6D). Histologically, UVB increased immune cell infiltrates compared to unexposed skin, forming immune aggregates primarily beneath dermal-epidermal junction and near hair sheath. Treatment with anifrolumab led to reduced immune infiltrates (Fig. S6B).

Transcriptionally, the transition from *LYVE*^+^ to *MMP9*^+^ *CD14*^+^ myeloid cells was evident in UVB-irradiated skin (Fig. 6E), with the *MMP9*^+^ population exhibiting a more proinflammatory phenotype. In non-lesional CLE skin, CD14^+^ cells were predominantly *LYVE*^+^ at baseline but shifted toward an *MMP9*^+^ state following UVB exposure, an effect that was markedly attenuated after anifrolumab treatment (Fig. 6F).

Spatially, UVB-induced *MMP9*^+^ *CD14*^+^ cells localized to the papillary dermis just beneath the dermal–epidermal junction or within discrete immune aggregates (Fig. 6G, S6C). Notably, anifrolumab treatment led to a substantial reduction in the accumulation of *MMP9*^+^ *CD14*^+^ cells at the dermal–epidermal junction, following the same dose of UVB, as confirmed by spatial transcriptomics (Fig. S6C).

UVB irradiation of non-lesional CLE skin also recapitulated several features characteristic of fibroblasts in lesional photosensitive skin, including thickening of the fibroblast layer at the dermal–epidermal interface and elevated expression of the chemokines *CCL2* and *CCL8*, both of which were attenuated following IFNAR blockade (Fig. 6H, S6D). Importantly, consistent with their migratory phenotype and mirroring findings from lesional CLE and DM in our scRNA-seq data, *MMP9*^+^ *CD14*^+^ cells expressed high levels of *CCR1* and showed slight upregulation of CCR5 and *CCR2*, the receptors for ligands including CCL2 and CCL8 (p<0.01) (Fig. 6I).

Together, these findings demonstrate that type I interferon signaling drives the UVB-induced migration of inflammatory *MMP9*^+^ *CD14*^+^ myeloid cells toward the epidermis, a key early step in the pathogenesis of interface dermatitis in photosensitive skin. Importantly, therapeutic blockade of IFNAR effectively suppresses this immunopathogenic cascade, supporting IFN-I pathway inhibition as a targeted strategy to prevent UVB-induced disease recurrence and exacerbation in photosensitive diseases.

## Type I interferon primes UVB-damaged keratinocytes to drive monocyte derived DC (moDC) activation and recapitulate MMP9**^+^** CD14**^+^** myeloid programs in photosensitive skin

Building on our observation that UVB triggers epidermal recruitment of inflammatory CD14^+^ myeloid cells in an IFN-I–dependent manner, we next investigated the role of keratinocytes, epithelial sentinels that absorb the majority of UVB radiation and initiate immune responses in photosensitive skin.

To this end, we used a keratinocyte–DC co-culture system designed to mimic key features of the lesional microenvironment. N/TERT2G immortalized keratinocytes exposed to UVB (50 or 100 mJ/cm^2^), with or without IFN-β priming, and their conditioned media were applied to CD14^+^ monocyte-derived dendritic cells (moDCs) generated from healthy donors (Fig. 7A). We selected moDCs ^8, 10^ as a surrogate for the MMP9^+^ CD14^+^ myeloid cells observed in CLE and DM lesions, given their shared expression of key markers *ITGAX (CD11c)*, *CD209*, *MAFB*^8^ (Fig. 2B, S2A).

**Figure 7.**
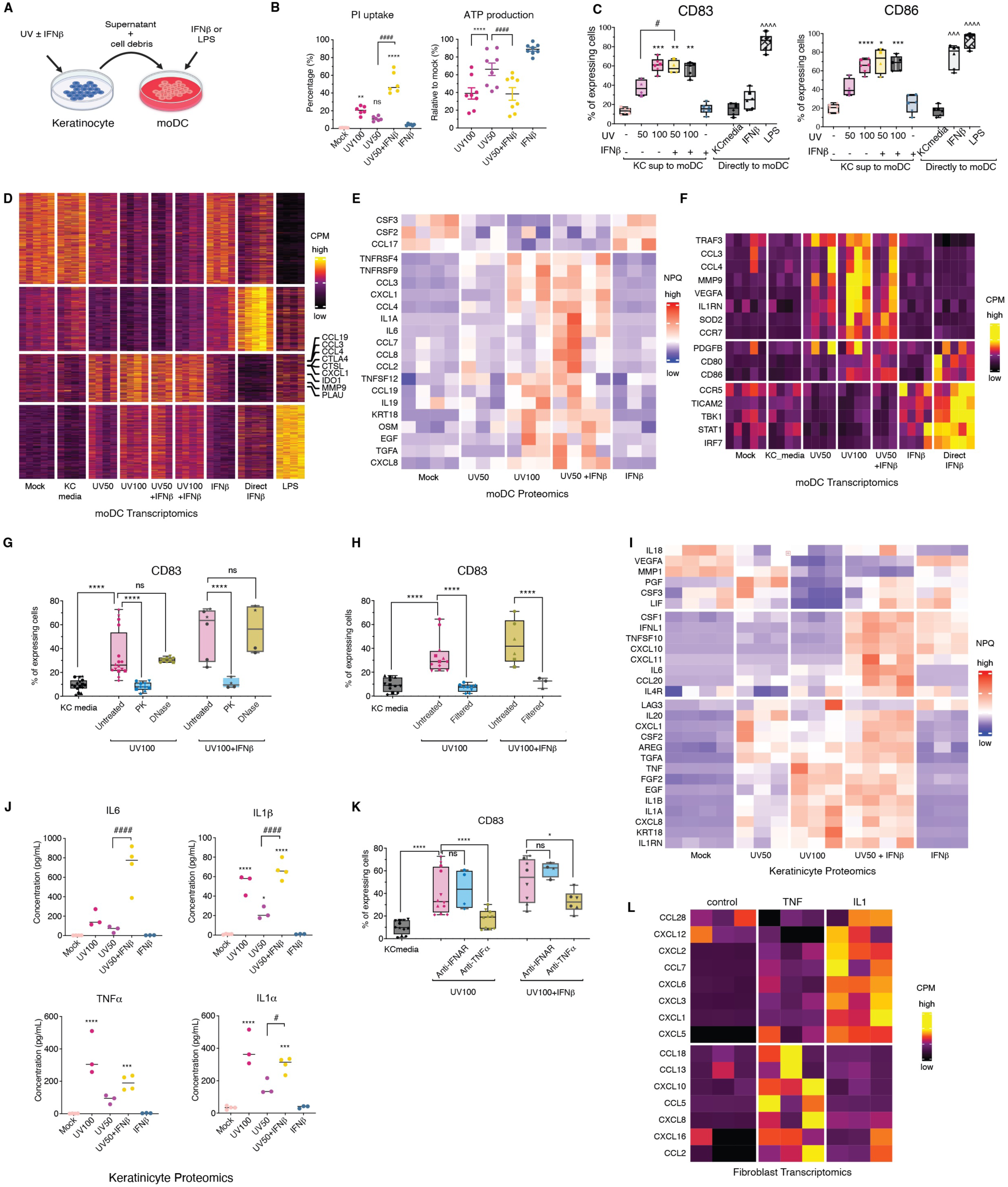
IFN-β–primed, UVB-irradiated keratinocytes activate and recruit inflammatory moDCs via proteinaceous, membrane-bound cytokines that trigger fibroblast-derived chemokine production. **(A)** Keratinocyte–moDC co-culture system. N/TERT2G keratinocytes were treated with UVB (50 or 100 mJ/cm^2^) ± IFN-β, and their supernatant were applied to monocyte-derived dendritic cells (moDCs). **(B)** Keratinocyte viability, measured by propidium iodide (PI) uptake and ATP depletion. Data are presented as mean ± standard error of the mean (SEM) and were analyzed using one-way ANOVA followed by Bonferroni-corrected pairwise comparisons. A p-value < 0.05 was considered statistically significant (n = 6–8) **(C)** Bar plots of moDC activation markers (CD83, CD86) expression after exposure to keratinocyte-conditioned media or direct IFN-β; lipopolysaccharide (LPS) stimulation as a positive control. Data are presented as mean ± SEM from five biological replicates (n = 4-6) and were analyzed using one-way ANOVA followed by Bonferroni-corrected pairwise comparisons. **(D)** Heatmap of differentially expressed genes (p < 0.05, |log_2_FC| > 1) in bulk RNA-seq of moDCs incubated with keratinocyte supernatants treated with UVB and IFN-β; data are are counts per million (cpm) scaled by gene (row). **(E)** Heatmap of differentially abundant proteins expression (NPQ) (p < 0.05) in moDCs after incubation with keratinocyte supernatants from the indicated conditions; colors scaled by protein (row). **(F)** Heatmap of MMP9^+^ CD14^+^ signature genes induced in moDCs after exposure to keratinocyte-conditioned media or direct IFN-β. Signature genes are M1-, M2-markers in Figure 5C that are high on the MMP9 end; data are counts per million (cpm) scaled by gene (row). **(G,H)** Percentage of moDCs expressing CD83 from flow cytometry. Data are presented as mean ± SEM and were analyzed using one-way ANOVA followed by followed by Bonferroni-corrected pairwise comparisons. **(I)** Heatmap showing significantly induced cytokines (p < 0.05) in keratinocyte supernatants under the indicated UVB and IFN-β conditions; colors are NPQ scaled by protein (row). **(J)** Cytokine concentration in keratinocyte supernatants, measured using OLINK proximity extension assay. Data are presented as mean ± SEM and were analyzed using one-way ANOVA followed by Tukey’s multiple comparison test . **(K)** Percentage of moDCs expressing CD83 from flow cytometry after incubation with keratinocyte supernatants pre-treated with TNF-α neutralizing antibody (adalimumab) or IFNAR blocker (anifrolumab) under UVB and IFN-β conditions. Data are presented as mean ± SEM and were analyzed using one-way ANOVA followed by followed by Bonferroni-corrected pairwise comparisons. **(L)** Heatmap of differentially expressed chemokines (p < 0.05, |log_2_FC|>1) in fibroblasts treated with IL-1α or TNF-α; data are counts per million (cpm) scaled by gene (row). (ns: not significant, * or # or ^ p < 0.05; ** or ## or ^^ p < 0.01, *** or ### or ^^^ p < 0.001, **** or #### or ^^^^ p < 0.0001).

UVB exposure increased keratinocyte membrane permeability (as measured by propidium iodide uptake; p < 0.01) and induced a dose-dependent reduction in intracellular ATP levels (p < 0.0001), both of which were further exacerbated by IFN-β pretreatment (p < 0.0001; Fig. 7B), indicating amplified epithelial stress and cell death. Conditioned media from UVB-treated keratinocytes robustly activated moDCs with upregulation of surface expression of HLA-DR, CD83 and costimulatory molecules CD80 and CD86, and this activation was enhanced by IFN-β priming of the keratinocytes prior to UVB exposure, while direct addition of IFN-β to moDCs did not have this effect (Fig. 7C; S7A).

To characterize this response, we performed transcriptomic and proteomic profiling of moDCs across all conditions. UVB alone induced a transcriptional program that was dose-dependent (UV50 < UV100), including upregulation of inflammatory mediators (CXCL1, CCL3, and CCL19 (Fig. 7D, E). Remarkably, the response to UVB 50 + IFN-β mirrored or exceeded that seen with UVB 100, underscoring the amplifying role of IFN-I in shaping keratinocyte output. Importantly, a subset of chemokines that selectively recruit lymphocytes and classical CD14^+^ monocytes, CCL2, CCL7, and CCL8, were only induced in moDCs exposed to media from IFN-β–primed, UVB-irradiated keratinocytes (Fig. S7B). These monocyte-attracting chemokines were not induced by UVB alone, even at high doses, indicating a qualitative shift in keratinocyte-derived signaling under IFN-I priming (Fig. S7B). Consistent with this, CCL2, and CCL8 were also among the most highly expressed chemokines in CD14^+^ cells found in lesional skin of DM and CLE patients (Fig. S7C), highlighting the relevance of this IFN-I–dependent axis in vivo.

We next asked whether this activation could be recapitulated by IFN-I directly. While direct IFN-β stimulation of moDCs was sufficient to upregulate canonical interferon-stimulated genes (*STAT1*, *IRF7*, *TBK1*), it failed to induce the full inflammatory program seen with keratinocyte-conditioned media. In contrast, moDCs treated with UVB + IFN-β–primed keratinocyte supernatants robustly upregulated key effectors of pathogenic myeloid programming, including *MMP9*, *PDGFB*, *CD80*, and *CD86*, mirroring the in vivo transcriptional profile of CD14^+^ MMP9^+^ cells found at the dermal–epidermal junction in CLE and DM lesions (Fig. 7F).

Together, these data reveal that dual inputs from type I interferons, direct signaling to moDCs, and indirect licensing of keratinocyte-derived inflammatory cues, are required to fully activate the MMP9^+^ CD14^+^ myeloid program. This synergy recapitulates the chemokine and matrix remodeling–rich signature observed in lesional skin of photosensitive diseases. Moreover, the presence of a robust IFN-I signature in non-lesional skin of CLE and DM patients suggests that even clinically unaffected skin is transcriptionally primed for exaggerated immune responses to UVB. This priming underlies the heightened recruitment and activation of inflammatory myeloid cells and supports IFNAR blockade as a rational therapeutic approach to disrupt this early pathogenic axis in photosensitive autoimmunity.

## IFN-I–licensed keratinocyte cytokines activate moDCs and induce fibroblast-derived chemokine programs in a multicellular inflammatory cascade

To define the molecular mechanisms by which type I interferons (IFN-I) license keratinocytes to activate moDCs, we first examined the biochemical nature of the activating factors released from UVB-damaged keratinocytes. Conditioned media from IFN-β–primed, UVB-irradiated keratinocytes robustly activated moDCs, as measured by CD83 expression (Fig. 7C). This effect was abrogated by proteinase K but not DNase I treatment, indicating that the active components are proteins (Fig. 7G). Moreover, their activity was retained after the supernatant was subjected to low-speed pellet but completely lost after filtration through a 0.22 μm membrane, suggesting a vesicle-associated or membrane-bound origin (Fig. 7H).

Multiplexed proteomic profiling of the keratinocyte supernatants identified 28 differentially expressed proteins (adjusted ANOVA p < 0.05), which clustered into UVB-induced and IFN-β–licensed groups (Fig. 7I). Several canonical inflammatory cytokines were robustly induced by UVB alone and further amplified by IFN-β priming, including TNF-α, IL-1α, IL-1β, CXCL1 and CXCL8. Strikingly, however, a distinct subset, including CCL20, CSF1 and most notably IL-6, was only highly induced in the presence of IFN-β priming (Fig. 7J, S7D). This pattern indicates that IFN-I does not merely amplify the UVB response but qualitatively reprograms keratinocyte cytokine output.

Given the biochemical profile of the activating factor, protein-based and filterable, we hypothesized that membrane-bound TNF-α could be a key effector. Indeed, TNF-α is robustly induced by UVB, further enhanced by IFN-β, and exists in both soluble and membrane-bound forms. Neutralization of TNF-β using adalimumab led to a 57% reduction in moDC activation (p < 0.001; Fig. 7K). However, when keratinocytes were pretreated with IFN-β prior to UVB irradiation, adalimumab only partially suppressed moDC activation (32% CD83^+^ vs. 17% without IFN-β), implicating additional IFN-β–primed cytokines in sustaining the response. Notably, direct IFNAR blockade on moDCs had no impact, indicating that IFN-I acts primarily on keratinocytes to license the production of proinflammatory mediators that act on bystander immune cells. Among these, IL-1α and IL-6 are compelling candidates: both are selectively induced in IFN-β–primed conditions and are well-established activators of myeloid cells.

Although keratinocyte-derived signals robustly activated moDCs, keratinocytes themselves failed to produce the chemokines necessary to attract classical monocytes, CCL2, CCL7, and CCL8, even after UVB + IFN-β stimulation. This observation prompted us to ask which cell type supplies the initial chemotactic cues that draw the first wave of MMP9^+^ CD14^+^ myeloid cells into otherwise quiescent skin. Spatial transcriptomics pinpointed a discrete subset of pro-inflammatory fibroblasts, rather than keratinocytes, as the dominant source of these monocyte-recruiting chemokines in photosensitive lesions (Fig. 5F, S5B). Positioned immediately beneath the dermal–epidermal junction, these fibroblasts colocalized with infiltrating *MMP9*^+^ *CD14*^+^ cells, which express the cognate receptors *CCR1*, *CCR2*, and *CCR5* (Fig. S3F, 5F). This spatial proximity and ligand–receptor pairing suggest that fibroblasts, not keratinocytes, initiate the chemokine gradient that invites the first influx of inflammatory monocytes after UVB injury.

To test whether keratinocyte-derived cytokines trigger chemokine production in fibroblasts, we stimulated primary human dermal fibroblasts with TNF-α and IL-1β, the two cytokines most strongly upregulated in IFN-β–primed keratinocytes. This treatment was sufficient to induce robust expression of *CCL2*, *CCL7*, and *CCL13* in fibroblasts (Fig. 7L), confirming a cytokine-mediated relay in which UVB-injured keratinocytes instruct neighboring fibroblasts to produce monocyte chemoattractants.

Together, these findings reveal a multicellular inflammatory circuit in photosensitive skin wherein IFN-I–primed keratinocytes secrete membrane-bound and vesicle-associated cytokines (e.g., TNF-α, IL-1α, IL-6) that activate MMP9^+^ moDCs, while fibroblasts respond to these epithelial signals by producing chemokines that selectively recruit CD14^+^ myeloid cells. This feedforward loop, amplified by type I interferons, is abrogated by IFNAR blockade, highlighting a mechanistic rationale for targeting the IFN-I pathway in photosensitive dermatoses.

## DISCUSSION

Photosensitivity is a hallmark of cutaneous lupus erythematosus (CLE) and dermatomyositis (DM), yet its mechanistic basis remains incompletely defined. Much of our understanding has historically derived from mouse models, including those for lupus. However, fundamental biological differences limit their translational relevance. Rodents are nocturnal, lack sun-adapted skin, and their epidermis is far thinner than that of humans, allowing UVB to penetrate to the dermis, unlike in human skin, where it is almost entirely absorbed by the epidermis^32, 33^. Moreover, human basal keratinocytes uniquely express NLRP1, which mediates UVB-induced pyroptosis via ZAKα/p38 signaling, a pathway absent in murine keratinocytes^34, 35^. These disparities underscore the need for human-focused studies to accurately model photosensitivity and UV-triggered inflammation.

Here, through a comprehensive analysis of human skin from CLE and DM patients, we demonstrate that type I interferons (IFN-I) represent a unifying immunological axis underpinning photosensitivity across distinct disease states. While the central role of IFN-I in CLE is well established, our findings extend this paradigm to DM. We identify CD14^+^ myeloid cells as key producers of IFN-β in DM, and to a lesser extent in CLE, highlighting their conserved role in driving IFN-I-mediated inflammation. These results align with recent studies linking cutaneous DM severity to the enrichment of CD14^+^ monocytes and macrophages in lesional skin^36^. Our work refines the cellular identity of these IFN-I sources in human disease. While prior studies in DM suggested IFN-β production by CD11c^+^HLA-DR^+^ cells^36^, limited marker resolution precluded definitive classification. Using high-resolution single-cell and spatial transcriptomic approaches, we now characterize these cells as CD14^+^CD1c⁻ inflammatory dendritic cells, a population distinct from classical DCs, and establish them as the principal source of IFN-β in DM skin.

A particularly novel contribution of our study is the identification of MMP9^+^ CD14^+^ cells as central effectors of UV-triggered inflammation in both CLE and DM. These cells span a transcriptional continuum from LYVE1^+^ “tissue-resident” macrophage-like states to MMP9^+^ “inflammatory migratory” phenotypes enriched near the dermal-epidermal junction. Spatially restricted to immune clusters in superficial dermis, MMP9^+^ cells express IL1B, TNF, and inflammatory chemokines, suggesting an M1-like profile. Functionally, MMP9 is a matrix-degrading enzyme that can disrupt basement membranes and facilitate cell migration. Its upregulation in CD14^+^ cells correlated with their ability to traffic from dermis to the epidermal interface, and indeed, only the MMP9^high^ subset was efficiently captured by our suction blister technique, implying these cells are actively migrating toward the epidermis. Previous studies observed MMP9 elevation in CLE skin and its correlation with disease activity^37^, but its cellular origin and mechanistic role were unknown. Our work fills this gap, identifying MMP9^+^ CD14^+^ cells as the key source and establishing MMP9 as both a marker and mediator of skin barrier disruption.

Notably, although IFN-I (particularly IFN-β) has been reported to suppress MMP9 transcription in vitro^38^, our findings reconcile this by suggesting an indirect in vivo effect: IFN-I licenses UV-damaged keratinocytes to release inflammatory cytokines (e.g., TNF-α, IL-6, IL-1α), which then drive CD14^+^ differentiation into MMP9-high states. Thus, IFN-I blockade does not directly repress MMP9 in myeloid cells but effectively prevents their emergence by halting upstream priming events.

A second dimension of our study is the discovery of cytotoxic CD4^+^ T cells in lesional CLE and DM. These cells co-express Th1 markers (*TBX21*, *CXCR3*) and cytotoxic effectors (*GZMB*, *GZMK*, *PRF1*)^24^, and are absent in healthy skin or other inflammatory conditions like psoriasis. We show that IFN-β selectively enhances granzyme B production in activated CD4^+^ T cells in vitro, suggesting that IFN-I not only recruits Th1 cells but also endows them with cytolytic capacity. This broadens the canonical view of IFN-I as a Th1-polarizing cytokine, implicating it in direct effector licensing of CD4^+^ T cells, particularly relevant in DM lesions, where IFN-I and GZMB^+^ CD4^+^ T cells were most abundant. These findings provide a cellular basis for the interface dermatitis seen in CLE and DM, potentially driven by both cytotoxic CD8^+^ and licensed CD4^+^ effector cells.

Mechanistically, our data support a feed-forward inflammatory circuit in photosensitive skin. Chronic IFN-I exposure preconditions keratinocytes, and subsequent UVB damage triggers the release of IL-1α, IL-6, and TNF-α, particularly in the context of prior IFN-priming. These cytokines activate dermal fibroblasts, which then produce chemokines that recruit CD14^+^ myeloid cells. MMP9 expression enables these cells to breach the basement membrane and reinforce inflammation through cytokine release and IFN-β production. Spatial transcriptomics and co-culture experiments map this cascade across cellular compartments, with fibroblasts and keratinocytes acting as amplifiers rather than bystanders. Membrane-bound TNF-α emerged as a key effector in this loop, activating monocytes and reinforcing their pathogenic differentiation. This explains how even a modest UVB exposure can elicit exaggerated inflammation in IFN-primed skin.

Our study extends prior observations of an IFN-I signature in non-lesional CLE skin to DM^39, 40^, suggesting that this is not a disease-specific phenomenon, but rather a shared characteristic and predisposing mediator of photosensitivity. MMP9^+^ CD14^+^ cells are largely absent in non-lesional skin of CLE and DM, suggesting that basal IFN-I expression arises from other sources. In lupus, prior studies reported elevated keratinocyte-derived IFN-κ in non-lesional skin^41, 42^, establishing a tonic IFN-I milieu that heightens UV responsiveness. Although similar studies in DM are limited, we observed comparable IFN-I signatures in non-lesional DM keratinocytes, supporting a parallel mechanism.

Beyond correlative associations, our photoprovocation trial with anifrolumab (anti–IFNAR) offers functional evidence that type I interferons serve as the upstream licensing signal for UV-induced exacerbation. IFN-I blockade nearly abolished the UVB-triggered recruitment of MMP9^+^ CD14^+^ myeloid cells, directly linking clinical response to interruption of this inflammatory axis. These findings align with recent clinical data demonstrating the robust efficacy of anifrolumab in CLE^43^ and parallel emerging reports of therapeutic benefit from IFN-β neutralization in the cutaneous manifestations of DM^44^. Together, these observations provide a mechanistic rationale for IFN-I–targeted therapies in photosensitive autoimmunity and identify MMP9^+^ monocyte-derived dendritic cells as a tractable, disease-relevant cellular target. Given their spatial proximity to the epidermis, these cells may also be amenable to local or topical therapeutic strategies.

Another key insight from our study comes from spatial transcriptomics, which uncovered a previously unappreciated role for dermal fibroblasts in orchestrating myeloid cell chemotaxis. Although keratinocyte-derived cytokines were sufficient to activate moDCs, keratinocytes themselves failed to produce classical monocyte-recruiting chemokines such as CCL2, CCL7, or CCL8. Instead, we identified a subset of superficial pro-inflammatory fibroblasts immediately beneath the dermal–epidermal junction as the major source of these chemokines. These fibroblasts expressed high levels of CCL2 and CCL8 and were spatially aligned with CCR2^+^ MMP9^+^ CD14^+^ cells, indicating a functional ligand–receptor relationship. This stromal population appears to act as a crucial intermediary, translating epithelial-derived cytokine signals (e.g., TNF-α, IL-1α) into a chemokine gradient that guides monocyte recruitment into the upper dermis. Our findings corroborate emerging evidence that fibroblasts in CLE are hyper⍰responsive to inflammatory stimuli, including IFNs and TNF, and produce chemokines that amplify skin inflammation^45^. Our functional co⍰culture experiments demonstrate that fibroblasts activated by IFN⍰licensed, UVB⍰injured keratinocyte cytokines (e.g., TNF⍰α, IL⍰1α) produce a robust chemokine cascade, directly linking keratinocyte damage to recruitment of inflammatory monocytes. Thus, fibroblasts emerge not merely as passive bystanders, but as essential drivers that spatially and temporally coordinate the early stages of UVB⍰induced inflammatory recruitment. This multi⍰cellular relay mechanism enriches the canonical model of interface dermatitis and suggests that therapeutic targeting of fibroblast⍰derived chemokines could effectively interrupt the initiation of photosensitive skin inflammation.

In summary, our study delineates a multicellular inflammatory cascade initiated by IFN-I-primed keratinocytes in response to UVB, leading to cytokine release, fibroblast activation, chemokine production, and the directed recruitment of MMP9^+^ CD14^+^ myeloid cells. These findings establish IFN-I not only as a driver of epithelial injury but also as a central coordinator of tissue-wide immune programming. The therapeutic relevance is highlighted by the observation that IFNAR blockade disrupts this entire cascade, providing a mechanistic rationale for targeting multiple nodes within this axis as potential disease-modifying strategies in photosensitive autoimmune skin disorders.

## Methods

### Study design

Study participants were recruited under an Institutional Review Board–approved protocol (H00021295, and H-14848) at the University of Massachusetts Medical School. Participants included individuals diagnosed with vitiligo, psoriasis, CLE, or DM based on a clinical exam performed by a dermatologist. Healthy individuals, matched by sex and age, were included as controls and were selected based on the absence of autoimmune or inflammatory skin diseases. The skin samples collected included healthy skin, lesional skin, and no lesional skin (unaffected, healthy-appearing tissue located at least 10 cm away from lesional areas). Patient samples were used for scRNA-seq, flow cytometry, proteomics and spatial transcriptomics as applicable (Table S1).

### Suction blister biopsies and processing

Suction blisters were collected following the procedure described in a previous publication^46^. Briefly, suction blisters were induced using a negative pressure of 10-15 mmHg at 40°C for 30-60 minutes. The blister fluid was collected using a syringe and centrifuged at 350g for 10 minutes at 4°C to separate the supernatant and cell pellet. The supernatant was flash-frozen for protein analysis, while the cell pellet was prepared for inDrop scRNA-seq and flow cytometry.

### InDrop scRNA-seq

Cells from blister fluid were resuspended and processed using the inDrop protocol developed by Zilionis et al.^47^, with a modification extending the unique molecular identifier (UMI) from 6 to 8 bases. Libraries were sequenced on Illumina NovaSeq and NextSeq platforms, producing raw FASTQ files. These files were processed through the inDrop processing pipeline [https://github.com/garber-lab/inDrop_Processing], which extracted reads with whitelisted barcodes, removed deduplicated reads using UMIs, aligned sequences to the GRCh38 genome using STAR^48^ , generated digital gene expression matrices using End Sequencing Analysis Toolkit (ESAT)^49^. ^48^ , generated digital gene expression matrices using End Sequencing Analysis Toolkit (ESAT)^49^. A custom GTF file was employed on the lab generated inDrop and bulk RNA-seq to merge gene annotations with overlapping 3’ ends. Detailed methods are available in the supplementary materials.

### scRNA-seq analysis

The UMI count tables were processed using Seurat v5^50^. Quality control retained cells with 200-4000 UMIs and fewer than 20 hemoglobin UMIs (HBA1, HBA2, HBB and HBD). Mitochondrial and ribosomal genes (starting with RPL or RPS) were excluded from the top 2000 highly variable genes prior to PCA. The top 20-30 principal components (PCs) were used for UMAP visualization and unsupervised clustering with functions FindNeighbors and FindClusters. Cells were first clustered into main cell types, followed by sub-clustering within each main type to group cells into finer clusters. Doublet clusters, identified by specific markers of other cell types, were removed. The remaining cells were then clustered again to identify main cell types and then further clustered lymphocyte cluster and myeloid cluster for finer subtypes.

For lymphocyte clusters, proliferating cells and interferon-responsive cells formed isolated clusters. To mitigate cell cycle effect,130 cell cycle-related genes were excluded from the PCA inputs. Interferon response of each cell was scored using 50 marker genes of the IFN-response cluster with the highest log2FC and FDR < 0.01, and then the effect was regressed out during scaling process before PCA. The visualizations were performed in ggplot2^51^. Cell type markers were visualized with DotPlot function in Seurat. Single-cell level gene expressions were visualized with VlnPlot.sc in AddOns package.

Differential expression (DE) analysis at main cell type level was performed using pseudo-bulk samples normalized to one million UMIs per sample, analyzed with DESeq2^52^. For finer subtypes, edgeR^53^ was adapted for the single-cell Seurat data structure^4, 54^ . ^53^ Statistical significance was defined as FDR < 0.05. Heatmaps were generated using the ComplexHeatmap package^55^ .

Single-cell compositional data analysis (scCODA)^56^ , a Bayesian framework that accounts for the dependencies inherent of proportion analysis, was applied to compare cell type proportions between lesional skin samples and healthy controls. Setting a higher expected FDR is encouraged to increase model sensitivity to include more cell types in the report. Multiple expected FDR levels were applied to detect the highly significant proportional changes, as well as report potential changes for more cell types. For the proportion heatmap, the percentage of cell types was first calculated in each disease and then scaled by cell types for visualization. PercentPlot.sc function in AddOns package was used for visualization of individual cell type proportions across multiple conditions.

Visualization functions are packed in R package AddOns available on github [https://github.com/garber-lab/AddOns].

### 10x scRNA-seq of three DM samples analysis

Raw count matrices were normalized with NormalizeData function LogNormalize method. with Seurat v5 package. Integrated with CCAIntegration method on top 30 PCs (using top 2000 variable genes selected with FindVariableFeatures function) with default parameters. UMAP and clustering were performed on the integrated top 30 PCs.

### GSE179633 scRNA-seq analysis

Count tables of all samples in GSE170633 were downloaded. Three samples presented issues: GSM6031365 (HCBIAO1117) and GSM6031370 (HC1119E) had empty barcodes.tsv.gz files, and GSM6031371 (HC1120E) contained a corrupted matrix file. To recover the two samples with missing barcodes, we generated pseudo barcodes.tsv.gz files. Ultimately, we successfully obtained data from 13 epidermal samples (3 healthy controls [HC], 5 DLE, and 5 SLE) and 16 dermal samples (4 HC, 5 DLE, and 7 SLE).

Each sample underwent standard quality control using the Seurat workflow. Doublets were identified using both scDblFinder and DoubletFinder. Given the large number of cells per sample, we opted for a conservative approach and removed any cells flagged as doublets by either tool. After QC and doublet removal, 162,773 dermal cells and 86,808 epidermal cells were retained.

Cell cycle effects were regressed out, and Harmony was applied to integrate dermal and epidermal samples into a unified manifold. This manifold was then subsetted and re-annotated into four major categories: Fibroblasts, Keratinocytes, Myeloid and Dendritic Cells (DCs), and a combined group of Schwann cells, Melanocytes, Endothelial cells, and Mast cells. These annotated subsets were then merged to form the final integrated manifold used for downstream analysis.

MC_CD14 cells were embedded and clustered separately. All samples of epidermis and dermis were combined. 500 top variable genes were selected with vst method for PCA. Top five PCs were used for integration with HarmonyIntegration method and k.weight = 1. 30. The integrated five dimensions were used to create diffusion map with DiffusionMap function in th destiny package ^57^ . The first two eigenvectors were used for diffusion map embedding visualization. The first diffusion component was used to order cells along the LYVE1 to MMP9 transition, the rank was used to score and color the cells. Heatmaps were generated with the ComplexHeatmap package. In heatmaps including the expression in MC_CD14 cells along the transition, cells were binned into 50 equal sized bins, and aggregated the average expression within the bins. Colors were scaled for each gene (row) across all bins and cell types together.

### Proteomic analysis

The NULISA™ (NUcleic acid Linked Immuno-Sandwich Assay) Inflammatory Panel (Table S2) quantified 250 proteins in blister fluid samples. Only samples with NULISA Protein Quantification (NPQ) values above the level of detection (LOD) were included in visualization and DE analysis.

Olink’s Proximity Extension Assay (PEA) - Olink® Target 48 Cytokine/Immune Surveillance Bundle (Table S4) measured absolute cytokine concentrations in cell culture media. Monocyte-derived dendritic cell (moDC) secretion levels were calculated by subtracting baseline protein levels in the added media from the final concentrations post-treatment.

Statistical analysis for NULISA data utilized t-test and linear mixed models. olink_ttest and olink_lmer from OlinkAnalyze package [<u>10.32614/CRAN.package.OlinkAnalyze</u>] were used for Olink. BoxPlot.nulisa function in AddOns package was used for visualization. Statistical test functions are packed in R package AddOns available on github [https://github.com/garber-lab/AddOns]

### In vivo UVB photoprovocation

Under IRB H00021295, we conducted UVB photoprovocation on eight healthy donors following a standardized photoprovocation protocol^58^. On day 1, threshold testing was performed on six areas of the lower back for UVB irradiation. UVB exposure was administered using a SolRx 100-Series Handheld device equipped with fluorescent bulbs (285–350 nm, Philips PL S 9W/12/2P, Solar System, Canada) at doses of 25, 50, 75, 100, 125, and 150 mJ/cm^2^.

On day 2, 24 hours after threshold testing, the minimal erythema dose (MED) for UVB was determined for each subject. These personalized MED values were used to calculate the UVB dose for photoprovocation sampling. For subjects whose 1.5 MED dose fell within the tested range (≤150 mJ/cm^2^), samples were collected 72 hours after the initial threshold testing without additional UVB exposure. However, if the 1.5 MED dose exceeded 150 mJ/cm^2^, subjects were asked to return for a separate session to receive the 1.5 MED dose, and samples were collected 72 hours after this second session.

Skin sampling from UVB-treated sites and matched untreated control sites were collected using suction blistering or or punch biopsy. In a subset of the study, UVB photoprovocation was also conducted before and after three monthly 300 mg intravenous doses of anifrolumab (administered every four weeks) in the same individual to assess treatment-associated changes in response to UVB exposure. Clinical skin disease activity was also assessed before and after treatment using the Cutaneous Lupus Erythematosus Disease Area and Severity Index (CLASI) ^59^.

### T Cell Isolation, activation, and cytokine treatment

CD4^+^ T cells were isolated from healthy donor peripheral blood mononuclear cells (PBMCs) using the CD4^+^ T Cell Isolation Kit (Miltenyi Biotec). The cells were resuspended in T cell culture media (RPMI supplemented with Hepes, penicillin-streptomycin, and fetal bovine serum) and plated in 24-well Nunc cell culture plates.

For T cell activation, wells were pre-coated overnight at 4°C with anti-CD3 (10 ng/mL, BioLegend, Ultra-LEAF™ Purified anti-human, Clone OKT3) and anti-CD28 (2 ng/mL, BioLegend, Ultra-LEAF™ Purified anti-human, Clone CD28.2) antibodies dissolved in PBS.

After plating, the cells were treated with cytokines—either IFN-β (50 ng/mL, R&D Systems) or IFN-γ (100 ng/mL, R&D Systems)—and cultured for three days. Following the incubation, cells were stained for viability and surface markers CD2 and CD4. Subsequently, the cells were fixed, permeabilized, and stained intracellularly for Granzyme B (GZMB, BioLegend, Clone QA16A02) before being analyzed by flow cytometry.

### Cytokilling assay

The cytokilling assay was performed by measuring impedance of the cells using the xCELLigence Real Time Cell Analysis technology (RTCA, Agilent, Santa Clara, CA). Twenty thousand BT-20 cells (ATCC, cat. No HTB-19) were seeded in a RTCA E-96 well plate and left to adhere for 6 h. Then, 2 × 105 freshly isolated CD4 T cells from PBMC were added. Solitomab (Sigma Alrich), a bispecific antibody anti-CD3 and anti-epithelial-cell-adhesion-molecule, was used to stimulate the cells at a concentration of 100 ng/ml. Interferon beta (R&D) at 10, 100 or 1000 ng/ml or IFNg (R&D) at 100 ng/ml were added to the culture. Cells were left in the RTCA analyzer until maxim cell death was reached (∼ 40 h). The percentage of cytolysis was calculated with the measurement of the impedance and using the software provided with the RTCA technology.

### FFPE Single-Cell RNA sequencing and data analysis

Three lesional biopsies from 3 patients with dermatomyositis were processed for single-cell RNA sequencing. For each sample, two formalin-fixed paraffin-embedded (FFPE) curls (25 μm each) were dissociated using the Miltenyi Biotech FFPE Tissue Dissociation Kit (CG000632 RevA, 10X Genomics). The resulting cell suspension was divided equally into four centrifuge tubes, each containing approximately 300,000 cells. Cells in each tube were hybridized with a unique Probe Barcode, as per the instructions in the "Chromium Fixed RNA Profiling Reagent Kits for Multiplexed Samples" user guide (CG000527, 10X Genomics).Post-hybridization, cells from the four tubes were washed, counted, and pooled in equal proportions. Approximately 40,000 cells from the pooled suspension were loaded onto a Chromium Q chip (PN-1000422, 10X Genomics). Sequencing libraries were prepared and sequenced on an Illumina NovaSeq platform using paired-end dual-indexing (28 cycles for Read 1, 10 cycles for i7, 10 cycles for i5, and 90 cycles for Read 2).The sequencing data was demultiplexed using bcl2fastq (Illumina). The resulting FASTQ files were processed with Cell Ranger using the multi pipeline and the GRCh38-2020-A reference genome. Data analysis was performed using the Seurat package. Quality control and preprocessing steps included doublet removal using scDblFinder and ambient RNA correction using SoupX. The data were then normalized, and variable features were identified.To correct for technical variation, we applied Harmony integration using the first 20 principal components. UMAP dimensionality reduction was performed using the Harmony-corrected embeddings. Clustering was conducted using the Leiden algorithm with a resolution parameter of 1.2. Cell type annotation was performed using established marker genes, resulting in the identification of 15 major cell populations: eccrine glands, B cells/plasma cells, apocrine glands, melanocytes, lymphoid cells, myeloid cells, smooth muscle cells, Schwann cells, pericytes, sebaceous glands, lymphatic cells, endothelial cells, adipocytes, fibroblasts, keratinocytes, and mast cells. For subsequent analyses, we focused specifically on the myeloid cell population.

### Keratinocyte and fibroblast isolation and culture

Pannus samples were obtained from UMass Memorial Hospital under protocol #H11731. The tissues were stored in 70% ethanol and sterile normal saline solution (0.9% NaCl) prior to processing. Four to five 4-mm punch biopsies were taken from each pannus sample and placed in a solution of dispase I (25 mg/mL, #04942086001, Roche). The samples were incubated for 30 minutes and then stored overnight at 4°C.

The following day, the epidermis was separated from the dermis using sterile forceps and transferred to a 15 mL Falcon tube containing 2.5 mL of 0.25% trypsin-EDTA (#25200056, Gibco). The samples were incubated at 37°C for 10–15 minutes with gentle agitation to disaggregate the cells. Trypsin was neutralized with DMEM containing 10% FBS. The cells were then resuspended in Keratinocyte Serum-Free Media (KSFM) supplemented with epidermal growth factor (EGF) and bovine pituitary extract (BPE) (#17005042, Gibco), and 1% penicillin/streptomycin (#15140122, Gibco). N/TERT2G cells were also cultured in KSFM supplemented with EGF and BPE. Cultures were fed every other day and passaged upon reaching sub-confluence, approximately every 5–7 days.

For fibroblast cultures, the dermis from pannus samples was incubated in 1 mL of collagenase (1 mg/mL, #9001-12-1, Sigma) at 37°C for 1 hour. Collagenase was neutralized with DMEM containing 10% FBS, and a single-cell suspension was prepared by gentle pipetting. The cells were centrifuged, and the pellet was plated in DMEM containing 10% FBS and 1% penicillin/streptomycin to culture fibroblasts. Cultures were fed every other day and passaged upon reaching sub-confluence, approximately every 5–7 days.

For experiments, primary and immortalized keratinocytes were plated at 2 × 10 cells/well and 0.8 × 10 cells/well, respectively, in a 12-well plate and incubated for 1–3 days at 37°C. Fibroblasts were plated at 10 cells/well in a 6-well plate and incubated for 1–3 days at 37°C. Cell plating concentrations for other plate sizes were scaled up or down based on the well area.

### Dendritic cell differentiation and stimulation

PBMCs were isolated from de-identified, healthy donor leukopaks (UMass Leukocyte Core), using density gradient centrifugation with Ficoll-Paque PLUS (GE Healthcare # 17-1440-02). The interface layer containing PBMCs was collected, washed twice with PBS-EDTA, and resuspended in RPMI 1640 medium. Monocytes were enriched through CD14^+^ magnetic bead-based positive selection (EasySep Human CD14 Positive Selection Kit II, #17858, Stemcell Technologies) following the manufacturer’s protocol.

Monocytes were seeded in tissue culture plates at a density of 1×10 cells/mL in RPMI 1640 medium supplemented with 10% fetal bovine serum (FBS), 1% penicillin-streptomycin, 2 mM L-glutamine, 50 ng/mL recombinant human GM-CSF (BioLegend, #572904), and 50 ng/mL recombinant human IL-4 (BioLegend, #574006). The cells were cultivated at 37 °C and 5% CO2. On day 3, half of the old medium was carefully removed and replaced with a fresh medium containing the same cytokine supplements. On day 7, cells were stimulated for 24 hours with either keratinocyte-conditioned media (generated by treating NTERT keratinocytes followed by UVB irradiation), ultrapure LPS (100 ng/mL, E. coli O111:B4, Invitrogen), recombinant human IFN-β (R&D Systems, #8499-IF-010/CF), or recombinant human TNF-α (50 ng/mL, BioLegend, #570102). For selected experiments, conditioned media were pretreated with Proteinase K (200 μg/mL; QIAGEN, #RP107B-1) in 50 mM Tris-HCl with 1 mM CaCl at 55°C for 1 hour, followed by heat inactivation at 95°C for 10 minutes. DNase I treatment (100 U/mL; QIAGEN, #79254) was performed at 37°C for 30 minutes in DNase buffer (10 mM Tris-HCl pH 7.5, 2.5 mM MgCl , 0.5 mM CaCl ). In additional experiments, media were filtered through a 0.22 μm filters (MilliporeSigma, #SLGVR33RS) to assess the size of active components. For blocking experiments, cells were pretreated with adalimumab (MedChemExpress, #HY-P9908, 20 μg/mL, 1 hour; anti-human TNF-α antibody) or anifrolumab (MedChemExpress, #HY-P99168, 50 μg/mL, 24 hours; anti–type I IFN receptor antibody) prior to stimulation. Following incubation, cells were harvested in Buffer RLT (Qiagen, #79216) for RNA extraction or processed for flow cytometry. Culture supernatants were collected for proteomic analysis.

### In vitro UVB treatment

For UVB irradiation, 0.8 -1× 10 N/TERT2G cells were seeded in 12-well clear plates (#150628, Nunc™) three days prior to irradiation. The cells were washed once with phosphate-buffered saline (PBS, pH 7.4) before being exposed to the indicated dose of UVB (50 mj/cm^2^ or 100 mj/cm^2^) using a SolRx 100-Series Handheld device equipped with fluorescent bulbs (285–350 nm, Philips PL S 9W/12/2P, Solar System, Canada). After UVB exposure, the PBS was replaced with 500 μL of keratinocyte medium, and the cells were incubated at 37°C with 5% CO for 24 hours.

### Membrane permeability assays

The cells were incubated with media containing Hoechst 33342 (1:30,000; Invitrogen, #P3566) and propidium iodide (PI, 1:1500; Invitrogen, #H3570). Imaging was conducted every 2 hours for a total of 24 hours using a Lionheart FX automated microscope (Biotek). Images were analyzed with Gen5 software, and the ratio of PI-positive to Hoechst-positive cells was calculated to determine the percentage of cells with compromised membrane integrity.

### ATP production assay

ATP levels in keratinocytes were quantified using the CellTiter-Glo 2.0 Assay (Promega, #G9242), following the manufacturer’s protocol. Briefly, keratinocytes were cultured in 12-well plates at appropriate densities and treated under experimental conditions. At 24 hours, an equal volume of CellTiter-Glo 2.0 reagent was added to each well, and the plate was incubated at room temperature for 10 minutes to allow for complete cell lysis and stabilization of the luminescent signal. Luminescence, proportional toethe ATP content, was measured using a microplate reader. The resulting luminescence values were normalized to control samples and used as a measure of cellular metabolic activity.

### Flow cytometry

The cells obtained through suction blister skin sampling were treated with Fc receptor blocking solution Human TruStain FcX (Biolegend, #422302) and stained with the following dye and fluorescent antibodies according to the manufacturer’s instructions: Zombie NIR™ Fixable Viability Kit (Biolegend, #423106), PerCP anti-human CD45 (Biolegend, #368506), Brilliant Violet 510™ anti-human CD3 (Biolegend, #344828), Spark NIR™ 685 anti-human CD19 (Biolegend, #302270), BUV737 anti-Human CD56 (BD Biosciences, #612766), Brilliant Violet™ 570 anti-human CD16 (Biolegend, #302036), Spark Blue™ 550 anti-human CD14 (Biolegend, #367148), PE/Fire™ 810 anti-human HLA-DR (Biolegend, #307683), eFluor™ 450 anti-human CD11c ( eBioscience™ #48011642). Cultured DCs were detached using ice cold PBS, and prepared and stained with Fc receptor blocking solution, Live dead zombie green (Biolegend #423111), CD11b (Biolegend, #101226), CD11c (Biolegend, #301608), CD14, HLA-DR (BD #562804) , CD83 (BD 612823), CD86 (Biolegend, #305442) and CD80 (BD #563315). An Aurora cytometer from Cytex Biosciences was used for sample analysis, and FlowJo™ Software (Version 10.9.0) for data processing.

### Bulk RNA-seq

Total RNA was extracted from frozen samples in RLT buffer using the RNeasy Mini Plus Kit (QIAGEN). To eliminate residual genomic DNA, the RNA was treated with RNase-free DNase I for 15 minutes at room temperature. RNA-seq libraries were constructed using an optimized CellSeq2 protocol for bulk RNA libraries (Gellatly et al., 2021). Each library was prepared by pooling 24 samples, with 50 ng of input RNA per sample.

The RNA libraries were sequenced on the NovaSeq 6000 platform. Raw FASTQ files were processed using the following pipeline (https://github.com/shuo-shan/viafoundry-celseq2-bulk-pipeline), with a modified GRCh38 genome as the reference (details in Supplementary Material), to generate the final count matrices, that were then analyzed in R (version 4.3.3).

### Public data acquisition

The data used for the Dendric cells stimulated with IFNγ analyses in this paper come from publicly available repository. We obtained the data from GEO: **GSE268354.** Raw FASTQ files were processed using the following pipeline (https://github.com/Carol-Salomao/RNA-seq), with a modified GRCh38 genome as the reference (details in Supplementary Material). After alignment, the data were analyzed using R. Data from Zheng et al.^22^ (GEO: **GSE179633**) was in the format of raw UMI count tables. Samples were integrated with Harmony, and then clustered and analyzed with Seurat v5 ^50^and AddOns packages.

### Spatial genomics gene positioning system (GenePS)

A custom seqFISH panel containing 521 genes, including 21 genes to be imaged one-at-a-time, was designed with Spatial Genomics (Primary Probe Kit, v500; No. 30021131). Fresh frozen human skin biopsy samples obtained at UMass Memorial Hospital under protocol (H00021295) were cryo-sectioned at 10 µm thick and mounted on specialized coverslips (Spatial Genomics, Catalog No. 10200003). Immediately after collection, the sections were fixed with freshly prepared 4% paraformaldehyde (PFA; Thermo Scientific, Catalog No. 28908) in 1× phosphate-buffered saline (PBS) for 15 minutes at room temperature. Following fixation, the sections were washed three times with 1× PBS for 5 minutes each and dehydrated in 70% ethanol for 30 seconds at room temperature. After air drying, the sections were stored at −80°C until further preparation for seqFISH experiments. Tissue processing was performed in accordance with Spatial Genomics’ Sample Processing for RNA Profiling Guide (ASY-053).

Tissue sections were imaged using the Gene Positioning System (GenePS; Spatial Genomics, Inc.), an automated platform for image acquisition, reagent delivery, and data processing. The system was loaded with the sample, decode plate, and buffers following the GenePS Instrument User Guide (INS-205) and guided on-screen instructions. Regions of interest (ROIs) within human skin sections were selected using DAPI-stained nuclei imaged with a 10× air objective. Once ROIs were selected, the automated imaging workflow was initiated, employing a 60× oil objective with standard image acquisition settings and capturing either three or six z-planes at a z-step size of 1.5 µm. The experimental protocol included iterative rounds of decode probe hybridization, imaging, and signal removal, continuing until all hybridization rounds were completed.

Raw image files were aligned across hybridization rounds and processed directly on the GenePS instrument to localize RNA signals. Subsequent analysis using Spatial Genomics’ Navigator software enabled transcript decoding and cell segmentation. The segmentation algorithm employed machine learning to delineate nuclei based on DAPI staining, with segmentation boundaries expanded by 15 pixels (∼1.5 µm) beyond the nuclei borders. RNA transcripts were decoded with a max allowed false discovery rate (FDR) of 5% and the transcripts were assigned to individual cells, generating cell-by-gene count matrices for each ROI. The processed outputs included a cell-by-gene matrix, cell coordinates, transcript list, DAPI image, and segmentation mask, providing high-resolution spatial and transcriptional data for downstream analysis.

### seqFISH+ analysis

Cell-by-gene matrices, cell coordinates and transcript locations were used to create the Seurat ^50^ object. Cells with fewer than 50 detected transcripts detected were excluded. For cells located at the skin surface, where transcriptional activity decreases during keratinocyte differentiation, a more lenient threshold of 10 transcripts was applied. To integrate samples from different patients, we used the RPCA integration method implemented in the IntegrateLayersfunction. For initial dimension reduction and clustering, we employed 30 principal components (PCs) and a resolution of 0.2. For finer subtype identification, clusters corresponding to endothelial cells, fibroblasts, keratinocytes, myeloid cells, and lymphocytes were processed separately. This included PCA, CCA integration, UMAP, and clustering using the top 20-30 PCs. For keratinocytes, the top 200 features were used for dimension reduction and clustering. For other cell types, the top 400 features were selected, excluding marker genes of other cell types identified using the FindMarkers function with thresholds of |log2FC| > 2 and adjusted p-value < 0.01. For MC_CD14 subtypes, PCA and CCA integration were conducted using the top 100 highly variable genes. The top five PCs were used for UMAP visualization and clustering. Cell types were annotated based on known marker genes.

To order MC_CD14 cells along the LYVE1^+^ to MMP9^+^ transition, top 100 variable features were selected by FindVariableFeatures and were then used for PCA. The top five PCs were used for CCAIntegration and the following DiffusionMap from the destiny package with parameter k=200. The first diffusion component was used to order the CD14^+^ cells along the transition. For expression heatmap, cells were binned into 30 equal-sized bins, and expression were aggregated within each bin. Colors of each gene (row) were scaled with all bins and cell types together.

Spatial images were generated using software Spatial Genomics Navigator Lite. Previously described annotations were imported for visualization, with segmented cells color-coded by clusters. Gene transcripts of interest were displayed as colored dots within the section.

Differential expression analysis was performed using the Model-based Analysis of Single-cell Transcriptomics (MAST) framework^60^, wrapped by Seurat. Differential analysis along the CD14^+^ MCs transition axis was performed using tradeSeq^61^.

Colocalization and visualization functions are packed in scSpatial R package available on GitHub (https://github.com/garber-lab/scSpatial/). Colocalization strength of each query cell within a reference cell type was calculated using createField function (with sd=400 pixels) and the getValueInField with default parameters. The summation of colocalization strengths of all query cells were used to represent the colocalization score between two cell types. For colocalization heatmap, colocalization scores were scale so that the total score of all cells in the sample was 0, and the score of the query cell type itself was set to 100. Heatmap visualization and clustering were performed using ComplexHeatmap R package^55, 62^. Please see the supplementary materials and methods for more details on colocalization analysis. Gene expression trends along colocalization scores or distances were assessed by calculating rolling averages using the FeaturePlot.cont.rollmean function with window.prop = 1. The depth distribution of each cell type was visualized using the ggridges R package (Wilke C (2024). https://CRAN.R-project.org/package=ggridges).

To explore the spatial distribution of CD14^+^ cells in skin, using other cell types as indicators for skin structures, we used minimum distance to the closest basal keratinocyte to represent the distance of a given CD14^+^ cell to epidermis. CD4 and CD8 cells were used to quantify the distance to lymphocytes infiltrated immune aggregates. Contrary to the colocalization analysis where denser cells indicate a higher potential interaction between two cell subtypes, cell density of the indicator cell types here won’t affect the spatial location and thus not factored in. The relationship between CD14^+^ cells transition scores and spatial locations (represented by minimum distance) was visualized using ggplot2.

The chemokines transcripts expression levels around a given CD14^+^ cell was calculated using calcLocalExpression function in scSpatial package with parameter radius = 400 (41.2μm). Local expression of chemokines as a function of CD14^+^ cells transition score was visualized with geom_smooth function with parameters method = "loess", span = 0.2.

The smoothed expression level of TNF and IFNG over spatial embedding was visualized with ImageFeaturePlot.contour function in scSpatial package with parameter sigma = 400, threshold = 0.9, n_levels = 6.

## Acknowledgments

We thank the patients of J.E.H., M.Rashighi and R.A.V. for the donation of time and tissues. The University of Massachusetts Medical School Flow Cytometry Core. The University of Massachusetts Medical School Department of Dermatology’s clinical research team.

## Funding

This work was funded by the NIH P50AR080593 (to J.E.H. and M.G.), NIAID 5U01AI176310 (to J.E.H, M.G., M.R), King Trust, Bank of America Private Bank, Co-Trustees Fellowship (to K.A), Dermatology Foundation and Skin of Color Society to (N.H.). This work was funded in part from generous gifts from the Edward Fischer GVH Fund and supported by the Siegel Family (to M. Rosenbach). K.W. is supported by a NIH-NIAMS K08AR077037, a Burroughs Wellcome Fund Career Awards for Medical Scientists, and a Doris Duke Charitable Foundation Clinical Scientist Development Award, M. Rosenbach is supported by Priovant.

## Author contributions

Conceptualization: Y.W., K.A., J.E.H., M. Rashighi, M.G.; Methodology: Y.W., K.A., N.S.H., M.G.; Investigation: Y.W., K.A., N.S.H., C.S.L., C.H.L.E., N.M., L.W., K.S.A., K.W., K.F., S.G., M. Rosenbach, R.A.V., J.E.H.; Visualization: Y.W., K.A., N.S.H.; Funding acquisition: K.A., N.S.H., J.E.H., M. Rashighi, M.G.; Project administration: K.A., N.S.H.; Supervision: M. Rashighi, M.G.; Writing: Y.W., K.A., N.S.H., M. Rashighi, M.G.

## Declarations of interests

M.G. is a scientific founder of Via Scientific Inc. which develops analysis pipelines used in this work. J.E.H. is a scientific founder of Villaris Therapeutics Inc., which develops therapeutic treatments for vitiligo. J.E.H. is inventors on patent #62489191 “Diagnosis and Treatment of Vitiligo,” which covers targeting IL-15 and Trm for treatment of vitiligo. K.W. receives research support from Merck Sharp & Dohme and 10X Genomics. M Rosenbach receives research support from Honoraria from Merck, J&J, Priovant, Novartis.

## Data and materials availability

All data associated with this study are present in the paper or the Supplementary Materials. *Data is in the process to be deposited in the ARK portal as required by the funding agency*. Functions used for analysis of the scRNA-seq data and seqFISH data are available on GitHub in open-source packages: https://github.com/garber-lab/addons; https://github.com/garber-lab/scspatial. Data, code, and materials will be made available upon request.

---

**Figure S1.**
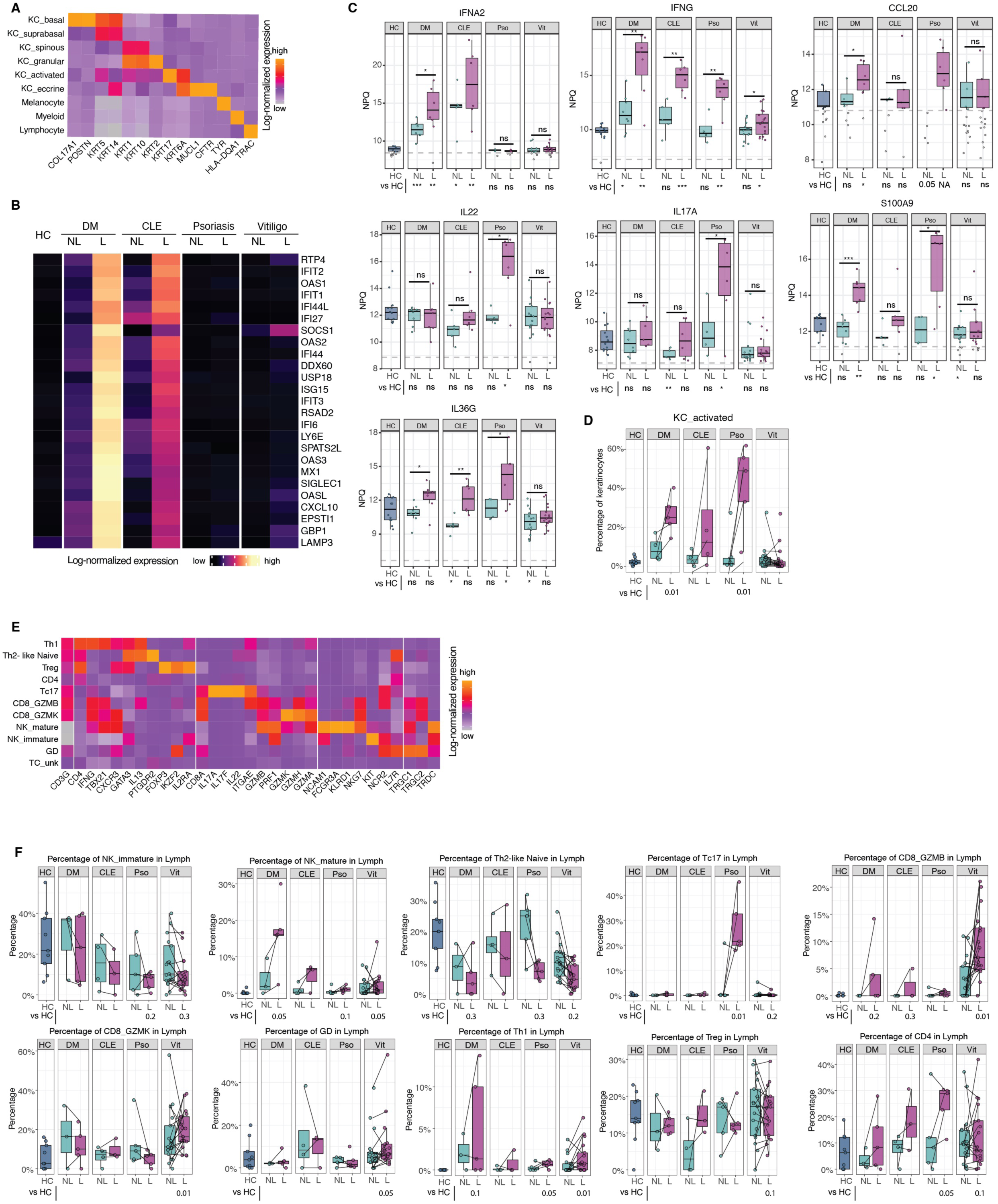
Keratinocyte and lymphocyte subtypes characterization and molecular signatures. **(A)** Pseudo bulk expression heatmap of marker genes for main cell types and keratinocyte subtypes in scRNA-seq of blister biopsies. **(B)** Pseudo bulk expression heatmap of type I interferon (IFN-I)–responsive genes in lesional (L) and non-lesional (NL) skin across diseases. **(C)** Box plots of protein levels (NPQ) in blister fluids across diseases and skins. T-test was used for pairwise comparisons: Not Significant (ns), \**P* < 0.05, ** *P* < 0.01, *** *P* < 0.001) **(D)** Box plots showing percentage of activated keratinocyte (KC_activated) within all keratinocytes (KC) per sample across skin conditions. Statistical significance was assessed using scCODA, with KC_suprabasal as the reference cell type. **(E)** Pseudo bulk expression heatmap of canonical marker genes for lymphocyte subsets. **(G)** Box plots showing percentage of lymphocyte subtypes per sample across conditions. Statistical significance was assessed using scCODA, with KC_suprabasal as the reference cell type.

**Figure S2.**
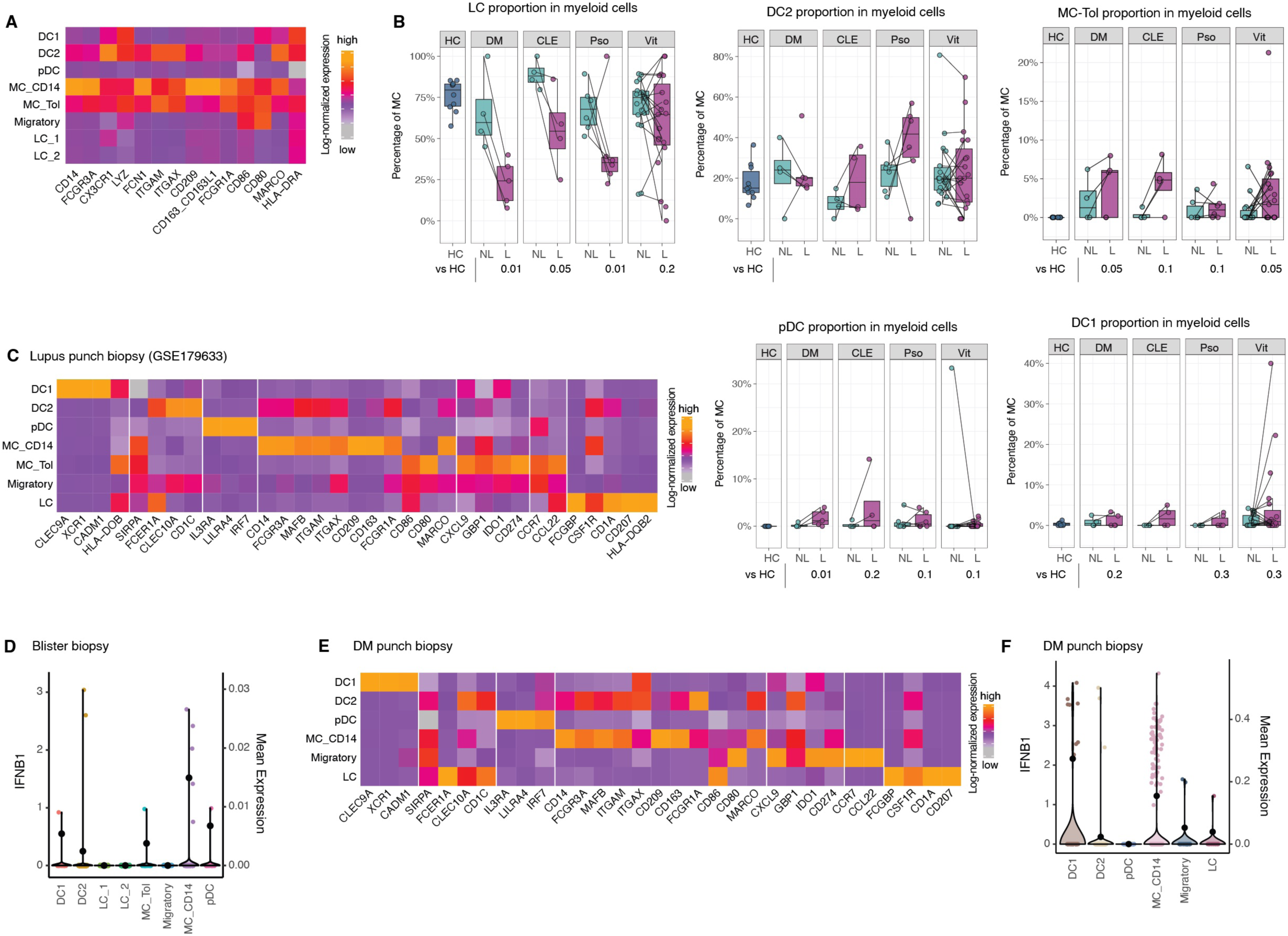
Cross-validation of myeloid subset composition and IFNB1 expression across scRNA-seq datasets in photosensitive skin diseases. **(A)** Pseudo bulk expression of macrophage and dendritic cell defining marker genes in scRNA-seq of blister biopsies. **(B)** Box plots showing percentage of myeloid subsets within all myeloid cells (MC) per sample, across skin conditions. Each dot represents a sample. Statistical significance was assessed using scCODA, with KC_suprabasal as the reference cell type. **(C)** Pseudo bulk expression heatmap of myeloid cell marker genes across myeloid subsets in punch biopsies obtained from lesional skin of lupus (GSE179633). **(D)** Violin plot showing the expression of IFNB1 from myeloid subsets detected in blister biopsies. Each dot represents a single cell. **(E)** Pseudo bulk expression heatmap of myeloid cell marker genes across myeloid subsets in scRNA-seq of punch biopsies obtained from lesional skin of DM. **(F)** Violin plot showing the expression of IFNB1 from myeloid subsets detected in DM punch biopsies. Each dot represents a single cell.

**Figure S3.**
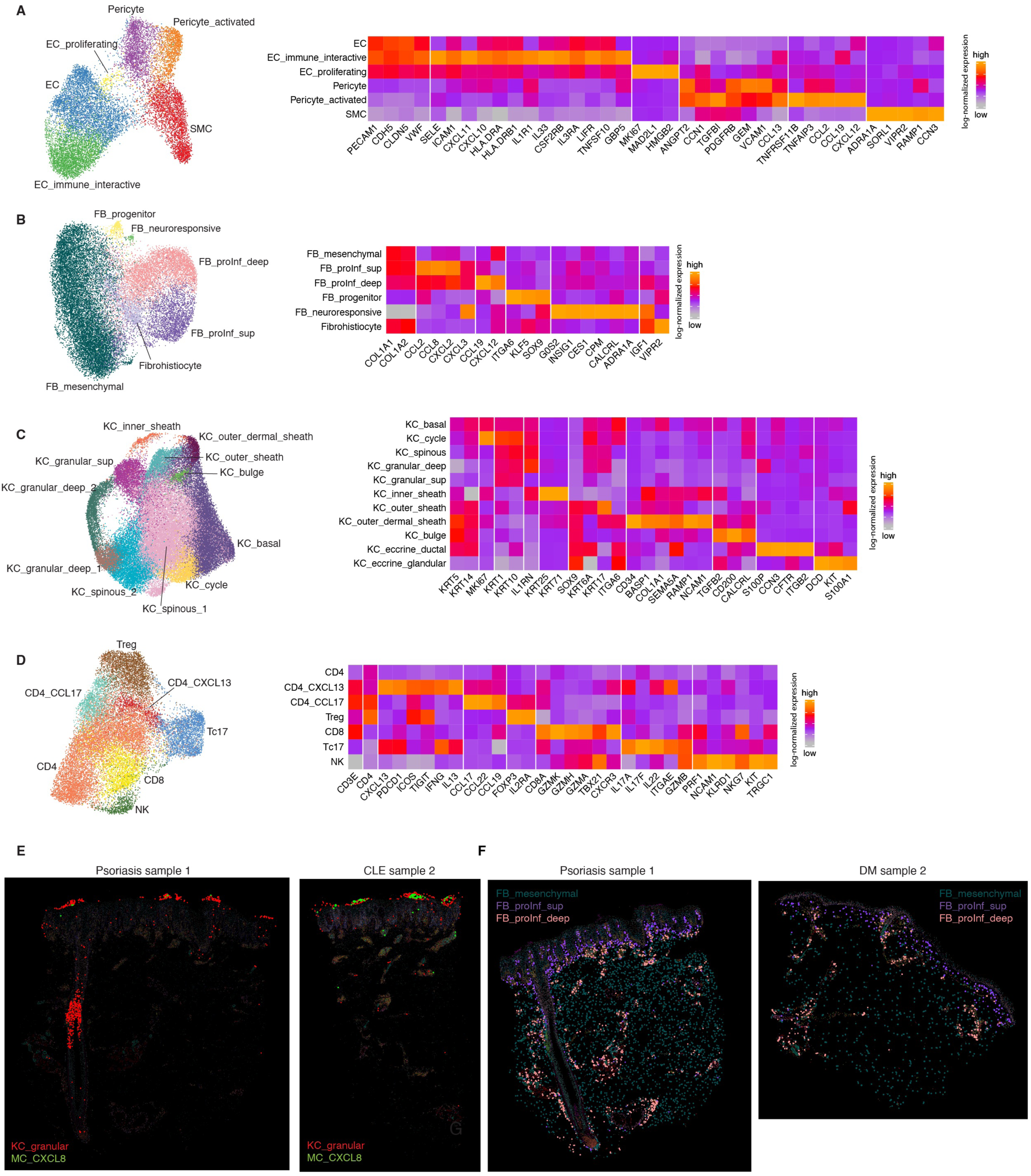
Spatial transcriptomics reveals fine-grained subtypes and spatial organization of stromal, immune, and inflammatory cell states in autoimmune skin lesions. **(A-D)** UMAP embeddings (left) and heatmaps for pseudo bulk marker genes expression of reclustered main cell types in seqFISH (A) Endothelial cells, (B) Fibroblasts, (C) Keratinocytes and (D) Lymphocytes. (E) Spatial embedding of seqFISH sections highlighting MC_CXCL8 cells using KC_granular as spatial reference. (F) Spatial embedding of seqFISH sections highlighting superficial proinflammatory fibroblasts (FB_proInf_sup), deep proinflammatory fibroblasts (FB_proInf_deep), and mesenchymal fibroblasts (FB_mesenchymal).

**Figure S4.**
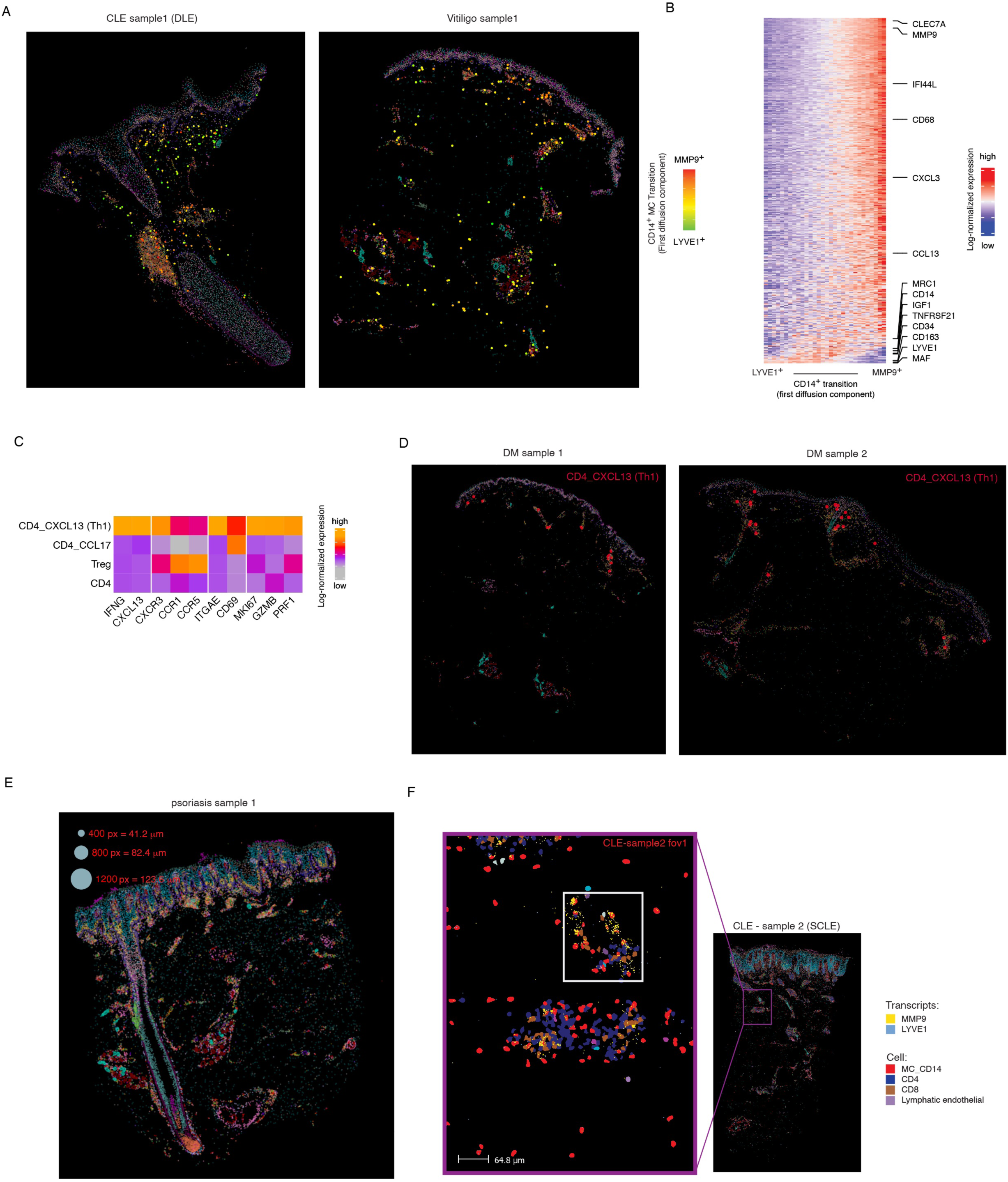
Spatial transcriptomics revealed transcriptional and spatial continuum along the LYVE1 to MMP9 transition. **(A)** seqFISH data showing CD14^+^ cells colored by their projection onto the first diffusion component in lesional tissue sections from the DLE and vitiligo samples. **(B)** Heatmap of differentially genes expressed in CD14^+^ cells ordered by the first diffusion component (transition) (p < 0.01, tradeSeq) and aggregated into 30 bins (193 cells per bin). **(C)** Pseudo bulk expression heatmap of cytotoxic and Th1 markers in different CD4 cell populations of the scRNA-seq data from blister biopsies. **(D)** Spatial embedding of seqFISH DM samples highlighting cytotoxic CD4_CXCL13 cell localization. **(E)** Representative spatial embedding illustrating radius used in colocalization analysis. **(F)** High-resolution image of CLE sample 2 showing zoom-ins of a dermal immune aggregate. Key cell types are displayed, with overlaid transcripts for LYVE1 (blue) and MMP9 (yellow).

**Figure S5.**
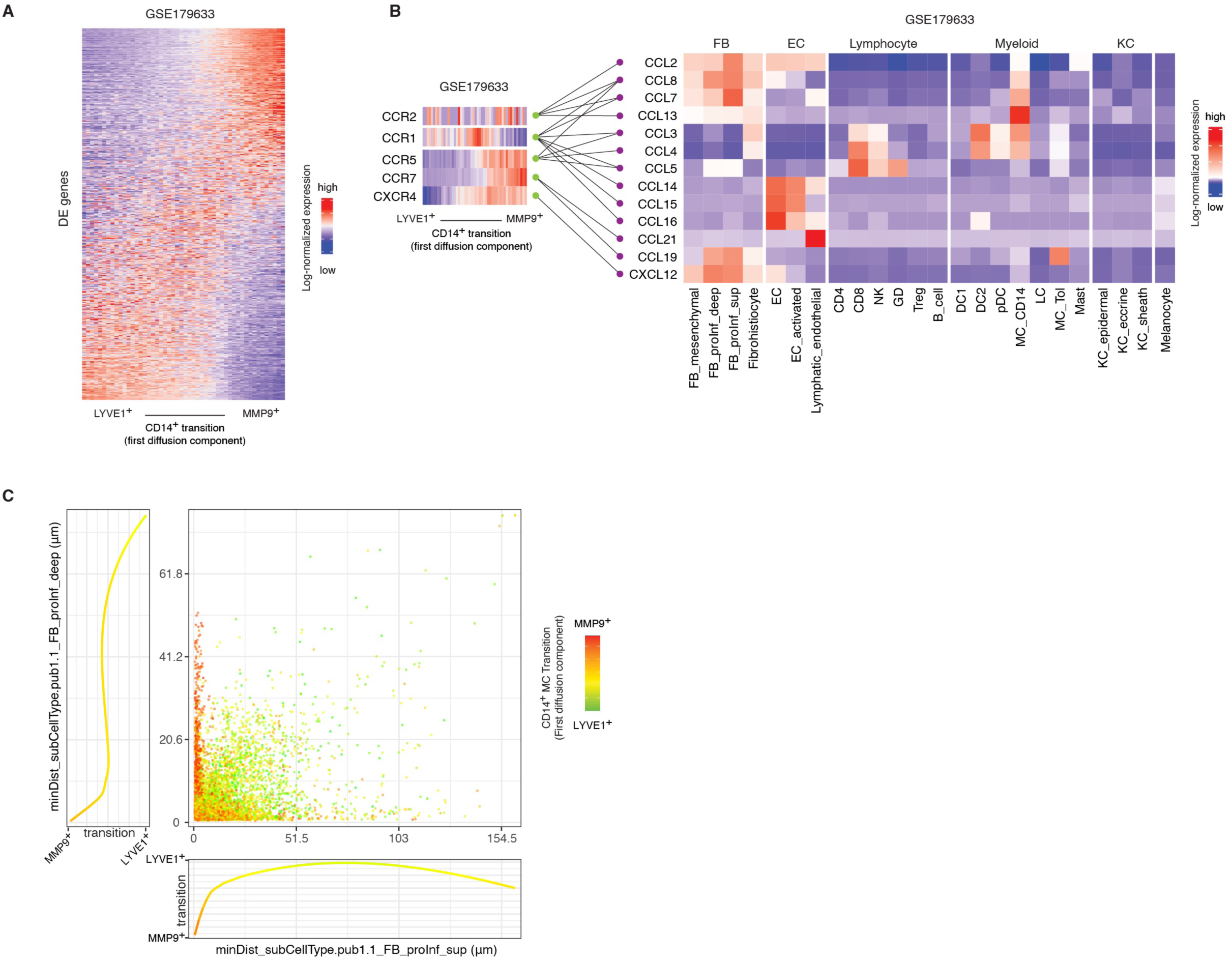
**(A)** Heatmap of differentially genes expressed across the CD14+ cell transition in the GSE179633 dataset (p < 0.01, tradeSeq). Cells were ordered by the first diffusion component (transition) and aggregated into 50 bins (144 cells per bin). **(B)** As in 5F, but with ligand pseudo bulk expression computed using the GSE179533 data. **(C)** Scatter plot showing minimum distance in μm from a CD14+ cell to a superficial pro inflammatory fibroblast (x-axis) or to a deep pro inflammatory fibroblast (y-axis) colored by their projection onto the first diffusion component; marginal plots show mean of the first diffusion component of CD14+ cells projected onto the x-axis (bottom) or the y-axis (left).

**Figure S6.**
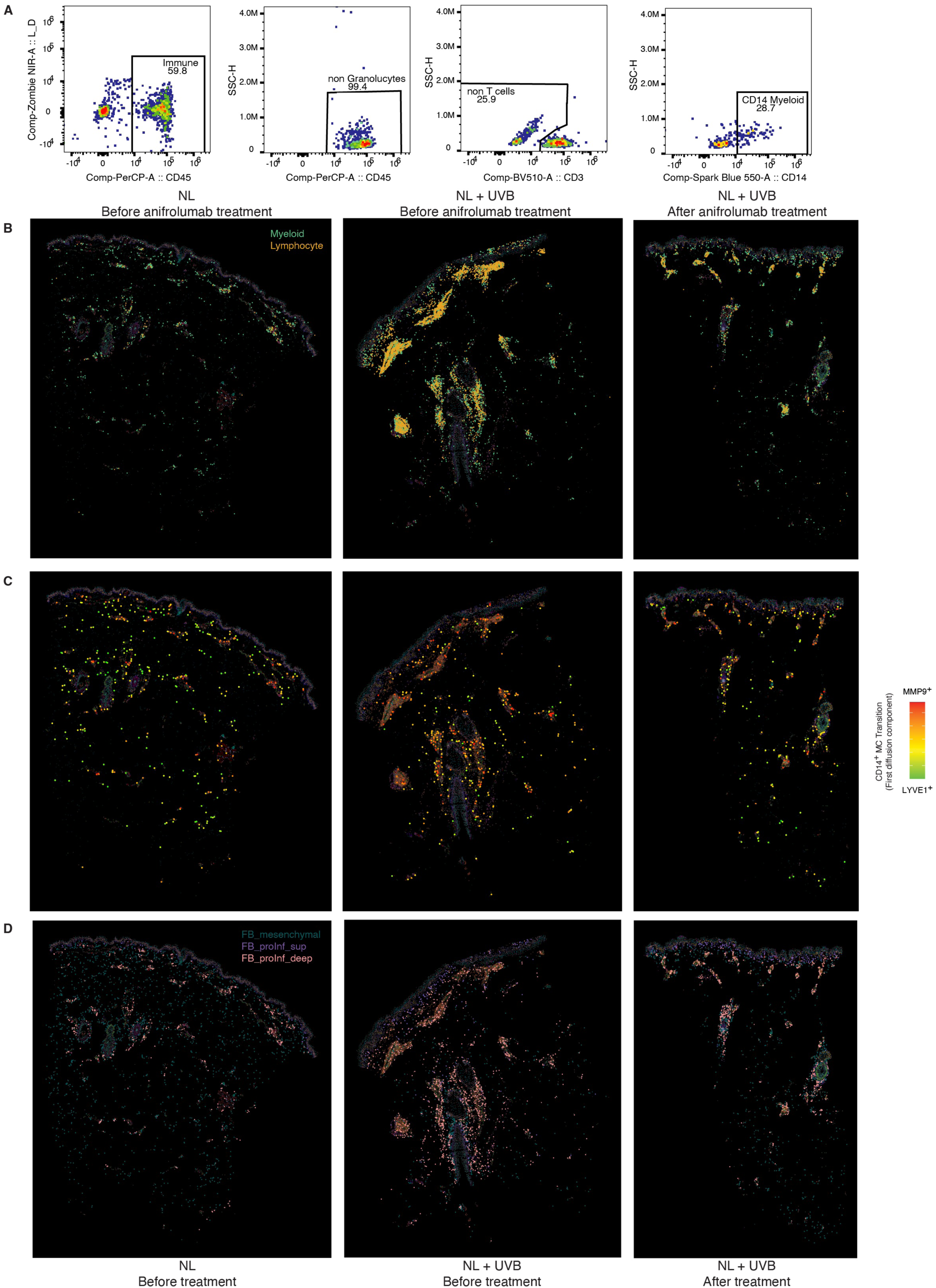
(**A**) FACS gating strategy to quantify CD14^+^ cells and myeloid cells in samples from non-lesional skin of a CLE patient. (**B-D**) Spatial embeddings highlighting relevant cell types in seqFISH data of the anifrolumab intervention experiment.

**Figure S7.**
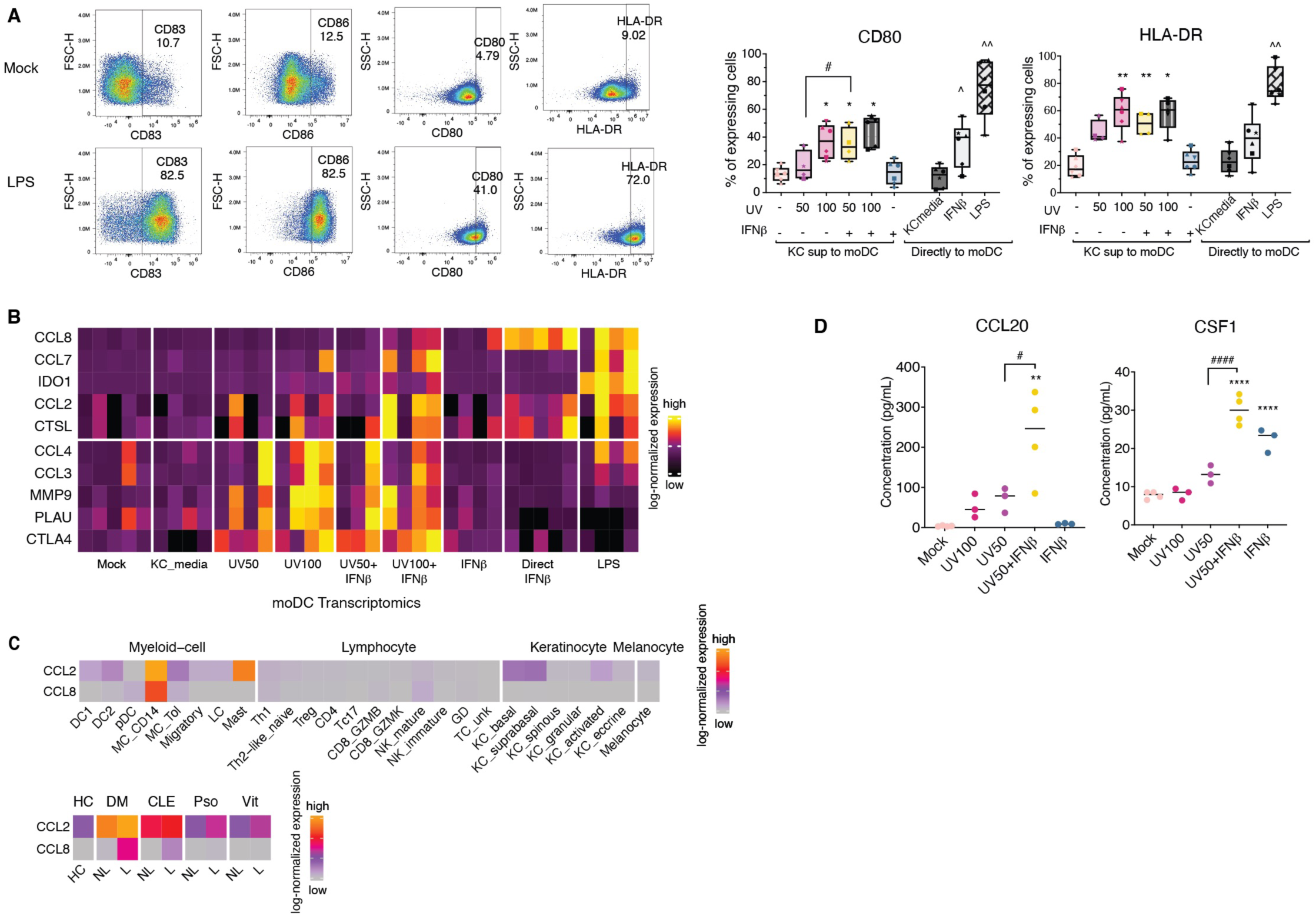
Type I interferon–licensed keratinocytes activate monocyte-derived dendritic cells (moDCs) and induce fibroblast-derived chemokine programs. **(A)** Left: Gating strategy for flow cytometric analysis of moDCs. Right: Quantification of CD80 and HLA-DR expression in moDCs incubated with keratinocyte supernatants or directly stimulated with interferon-β (IFN-β) or lipopolysaccharide (LPS). Data are presented as mean ± standard error of the mean (SEM) and were analyzed using one-way ANOVA followed by Bonferroni post hoc test. **(B)** Top: Heatmap of protein expression in moDCs incubated with keratinocyte supernatants under the indicated UVB and interferon-β (IFN-β) treatment conditions, or directly stimulated with IFN-β or lipopolysaccharide (LPS). Bottom: Quantification of selected proteins from the heatmap, shown as mean ± standard error of the mean (SEM). Statistical analysis was performed using one-way ANOVA followed by Tukey’s post hoc test. **(C)** Heatmap of selected inflammatory genes expression in bulk RNA-seq of moDCs stimulated with keratinocyte supernatants under the indicated UVB and interferon-β (IFN-β) conditions, or directly treated with IFN-β or lipopolysaccharide (LPS); colors are normalized expression scaled by gene (row). **(D) (H)** Cytokine Concentration in keratinocyte supernatants, measured using OLINK proximity extension assay. Data are presented as mean ± SEM and were analyzed using one-way ANOVA followed by Tukey’s multiple comparison test . (ns: not significant, * or # or ^ p < 0.05 ; ** or ## or ^^ p < 0.01, *** or ### or ^^^ p < 0.001, **** or #### or ^^^^ p < 0.0001).

## Notes

### Competing Interest Statement

The authors have declared no competing interest.

