## Supplementary Methods for "A Spatially Coordinated Keratinocyte-Fibroblast Circuit Recruits MMP9^+^ Myeloid Cells to Drive IFN-I-Driven Inflammation in Photosensitive Autoimmunity"

**Patient ID – sample name in the publication**

UV109 – DM sample1

UV253 – DM sample2

UV243 – CLE sample1

UV260 – CLE sample2

UV238 – Psoriasis sample1

UV239 – Psoriasis sample2

VB268 – Vitiligo sample1

**Supplementary Methods**

**Custom GTF file merging gene annotations with 3’ end overlap**

To address artifacts in single-cell RNA sequencing data, we created a custom GTF file that merges gene annotations with 3’ end overlaps.

Using the filtered gencode.v34.primary_assembly.annotation.gtf, as documented in 10x Genomics note for References 2020-A (July 7, 2020), we observed artificial clusters characterized by high and unique expression of 2-4 genes. These genes, such as DIABLO and AC048338.2, are neither recognized as cell type markers nor indicative of cell states. Closer inspection revealed that these artifacts arose from reads mapping to overlapping regions of adjacent genes, which were assigned to both genes by ESAT. The filtering of readthrough transcripts failed to resolve this issue.

To mitigate this while keeping the reads, we merged genes with overlapping regions within 800bp to the 3’ end, accounting for the 3’ capture bias inherent to inDrop technology. The merged genes are identified using meta IDs or by connecting gene names connected with underscores. Detailed merging records and nomenclature are available in the GitHub repository AddOns [<https://github.com/Yuqing66/AddOns/blob/main/data/metagene.txt>].

**Colocalization analysis**

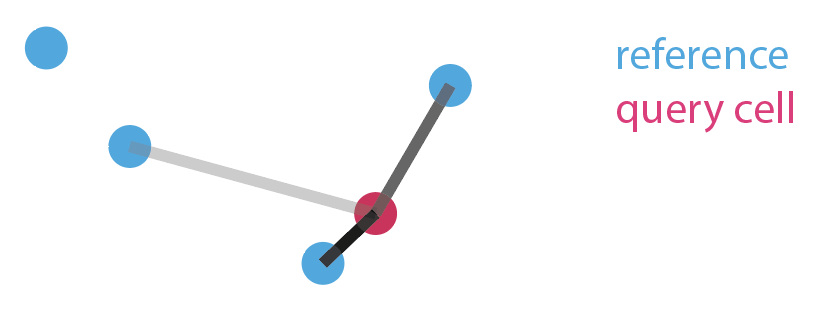

Cellular colocalization was quantified based on the number of neighboring cells and the distance between them. Connection edges were weighted by distance using a Gaussian transformation, with the edge darkness representing the weight in illustration. Adjusting the standard deviation of the Gaussian curve allows the analysis of interactions from direct contact to long-range cell-cell interactions, from refined to regional structures. The summation of all connection weights yielded the degree of colocalization for a given cell relative to a reference.

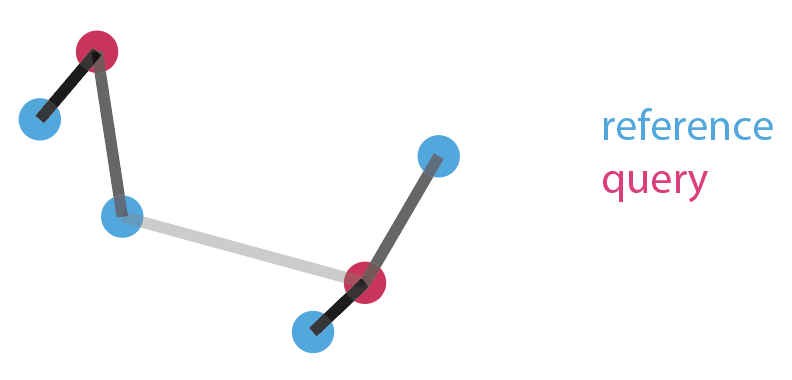

The degree of colocalization between two cell types was calculated as the sum of colocalization strengths of all query cells, normalized by the number of query cells. This metric indicates the extent to which query cells localize near the region of reference cells, but does not reflect reciprocal localization.

Additionally, we extended the analysis by weighing the Gaussian curve height with values of interest, such as the expression levels of specific genes. This approach is particularly useful when the localized gene expression was not captured by cell type clustering, such as keratinocyte activation and interferon responses. Furthermore, transcripts locations can server as input, enabling analysis of transcripts outside defined cell boundaries.
